# The geometry of cortical representations of touch in rodents

**DOI:** 10.1101/2021.02.11.430704

**Authors:** Ramon Nogueira, Chris C. Rodgers, Randy M. Bruno, Stefano Fusi

## Abstract

Neural responses are often highly heterogeneous non-linear functions of multiple task variables, a signature of a high-dimensional geometry of the neural representations. We studied the representational geometry in the somatosensory cortex of mice trained to report the curvature of objects using their whiskers. High-speed videos of the whisker movements revealed that the task can be solved by linearly integrating multiple whisker contacts over time. However, the neural activity in somatosensory cortex reflects a process of non-linear integration of spatio-temporal features of the sensory inputs. Although the responses at first appear disorganized, we could identify an interesting structure in the representational geometry: different whisker contacts are disentangled variables represented in approximately, but not fully, orthogonal subspaces of the neural activity space. The observed geometry allows linear readouts to perform a broad class of tasks of different complexities without compromising the ability to generalize to novel situations.

## Introduction

Making sense of the real world often requires the integration of sensory evidence across multiple sources of information. In some situations, this process involves only simple operations like linear summation. For example if we need to determine whether an object is close to our hand or not, we can just move our fingers until any of them touches the object [1, 2]. Summing the tactile feedback coming from all fingers and comparing it to a threshold would be sufficient to report whether the object was present or not. In other words, a linear decoder would be sufficient to perform this simple detection task. However, recognizing the shape of an object by touch could be a more challenging task, which might involve non-linear integration of the sensory inputs coming from multiple fingers.

Here we studied recent experimental data [3] to understand how mice perform a shape recognition task using their whiskers. Using high-speed videos of the whisker movements, we discovered that the task can be solved by simple linear integration of whisker features (linear decoder). A linear decoder is also the best predictor of the decisions of the animals. However, the neural representations in somatosensory cortex are better explained by a process of non-linear spatio-temporal integration. This type of non-linearity typically corresponds to high-dimensional representations in the neural activity space (G3; Fig. 1). This means that while the animal is performing the task, the points that correspond to the observed patterns of activity define a high-dimensional object. These representations confer flexibility because a downstream neuron can perform a multitude of different tasks by being able to separate the points of the object in many different groups. Despite the non-linearity of the neural responses, we also identified a low-dimensional structure in the geometry of the neural representations, reflecting some of the properties of abstract representations [4]. An abstract representation of two variables (e.g. contacts of whisker one, *C*_1_, and contacts of whisker three, *C*_3_) is shown in Fig. 1 (G1), where they form a low-dimensional geometry in the neural activity space. The points that correspond to the states of activation of a population of neurons now define a 2D square. These representations are called abstract because the variables are represented in approximately orthogonal subspaces (one axis for *C*_1_ and one axis for *C*_3_ in the figure), and thanks to this arrangement they have special generalization properties: the decoder trained to report the value of one variables is always the same no matter what the value of the other variables is. The observed representations are neither these abstract low-dimensional representations (G1), nor the less structured high-dimensional representations (G3), but they are closer to what is represented in G2. These representations constitute a non-trivial compromise between the flexibility of high-dimensional representations, which can be reused in a number of very diverse tasks, and the ability to generalize and robustness to noise of low-dimensional representations.

**Figure 1:**
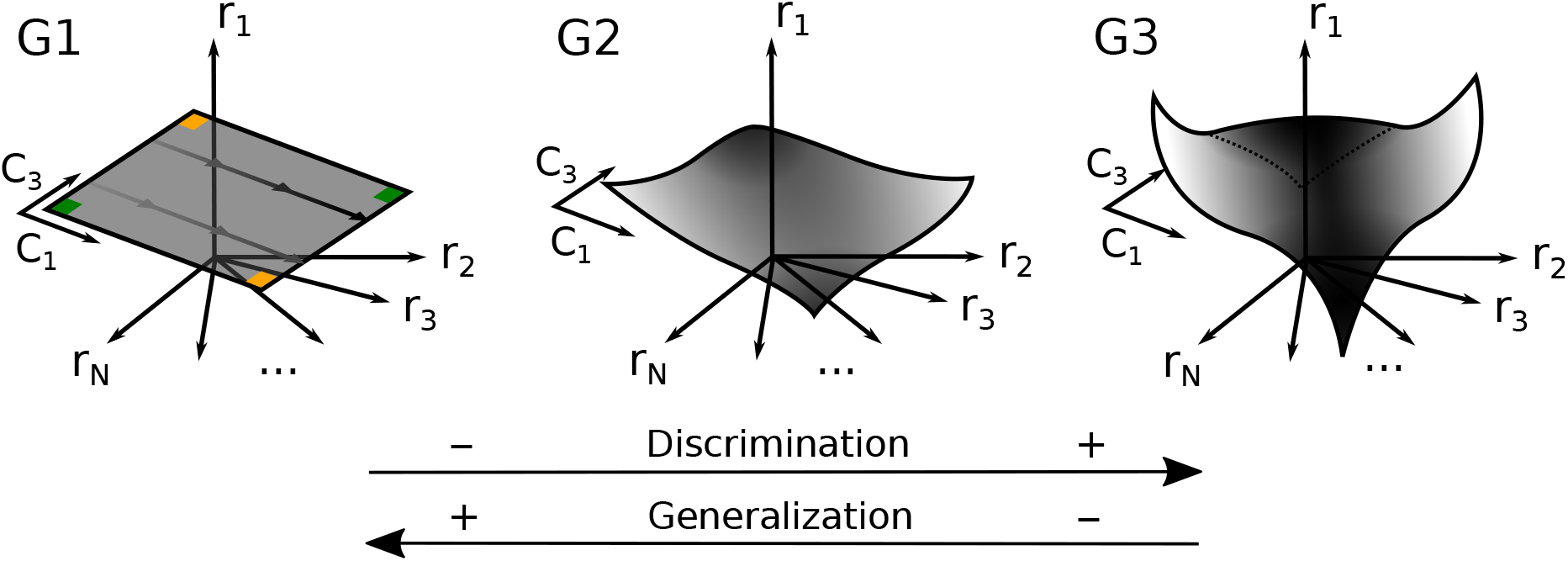
Different geometries of the neuronal representations are characterized by different computational properties. Each panel shows the firing rate space of *N* neurons, which is the space defined by the *N* orthogonal axes that represent the firing rate of the different neurons of the population. Each point in this space corresponds to one pattern of activity of the population of neurons. As the animal performs the task, the set of points visited is an object (gray) that has a particular geometry. Low-dimensional representations (G1): the number of contacts *C*_1_ and *C*_3_, two variables that we will show to be relevant for performing the task, are represented along two orthogonal axes. As one varies these two variables, the points that correspond to the neural activity define a 2D square. These representations generalize well to unseen experimental conditions (Generalization) but perform poorly on tasks that require non-linear combinations of different task variables (Discrimination). Indeed, a simple linear decoder trained to report the value of *C*_1_ (high vs low) would work for all possible values of *C*_3_ because the coding directions for *C*_1_ are the same for all values of *C*_3_ (parallel lines). So it could be trained only on one value of *C*_3_ and thanks to the low-dimensional geometry the decoder would work right away for a different value of *C*_3_. Analogously for *C*_3_: a decoder trained to report its value (high vs low) would work for any value of *C*_1_. This confers robustness to the responses of the readout, and allows for generalization in novel situations, e.g. when a *C*_1_ decoder is trained on one value of *C*_3_ and it is tested on a different value of *C*_3_, a condition never experienced by the decoder. The limit of these representations is that there are points that cannot be separated by a linear readout (a linear readout would be represented by a separating plane in these plots). For example the two points at the opposite vertices of the square cannot be separated from the other two opposite vertices (green vs orange). On the contrary, high-dimensional representations (G3) allow a simple linear readout like a neuron to separate the points on the gray object in a multitude of different ways. They are highly flexible but generalize poorly. Intermediate geometries (G2) could benefit from the computational properties of both low- and high-dimensional representations.

High-dimensional neuronal representations have been shown to enable a linear readout to perform a large number of different tasks [5, 6, 7]. This is a desirable feature in a brain area like prefrontal cortex, which is involved in many complex cognitive tasks. A linear readout of these high-dimensional representations (even by a single neuron) can be easily trained to generate a variety of response patterns, each corresponding to a different task. This can be achieved without modifying the input representations and confers flexibility to the neural system. The disadvantage of these high-dimensional representations is that the type of non-linear integration that they entail typically enhances noise and hence can impair the performance in simple tasks that depend on a linear combination of one or few variables (see e.g. [6]).

It is currently unclear whether the neural representations in sensory areas are high- or low-dimensional. Many experimental works have described the neural responses in sensory areas using the concept of a receptive field (see e.g. [8]). In particular, this was a successful approach in the pioneering work of Hubel and Wiesel in the primary visual cortex (V1) [9]. Receptive fields have also been derived for neurons in primary somatosensory cortex (S1), for example by reverse correlation of sparse noise stimuli applied to the whiskers [10]. The receptive field description of neuronal responses is useful only when the activity can be accurately predicted by a linear combination of input features, or in other words when the representations are low-dimensional. These representations guarantee robustness to noise, which is certainly a desirable property in a sensory area, but this typically happens at the cost of flexibility of being able to perform a broad range of tasks.

However, it is clear that even in brain areas like V1, there are significant deviations from linear models as soon as more complex, realistic stimuli are considered [11, 12, 13, 14]. In line with these observations, our study shows that the neural activity in somatosensory cortex of mice is best described by a non-linear model of the task variables. Although this is a sensory area, the neurons respond to rather heterogeneous non-linear combinations of these task variables, similarly to what has been observed in prefrontal cortex and in the hippocampus [15, 4]. This is compatible with the recent observations of high-dimensional representations in the visual cortex of non-behaving mice [16].

Heterogeneous non-linearities might suggest that the neural activity is rather disorganized, but in this case we could identify a low-dimensional scaffold in the geometry of the representations. This scaffold could enable downstream neurons to generalize to novel situations better than completely random and disorganized representations. In particular, the number of whisker contacts for different whiskers are represented along approximately orthogonal directions in the neural activity space. In other words, these variables are disentangled or factorized, and we know from the machine learning literature [17, 18] and recent experimental work [4, 19] that they allow for better generalization. This enhanced generalization induced us to consider these variables as abstract variables [4].

Our conclusion is that the geometry of the representations in somatosensory cortex is basically both low-dimensional and high-dimensional. It is low-dimensional because we can identify a lowdimensional structure that allows for enhanced generalization. At the same time, the non-linearities, which distort the low-dimensional scaffold, make the representations sufficiently high-dimensional to enable a downstream neuron to perform a broad class of complex discrimination tasks. The non-linear distortions are not so large that they compromise the robustness to noise and capacity to generalize. Geometry that balances complexity and generalization, as seen in prefrontal cortex and hippocampus of monkeys [4], may therefore be a feature of cortex in general. Our results suggest that it is important to use non-linear models to describe the neural activity in somatosensory cortex, but at the same time it is essential to analyze the neural activity at the population level to identify the organization of the neural activity space.

## Results

### The whisker-based object discrimination task

Mice were trained on a whisker-based shape discrimination task (Fig. 2a), in which they were asked to identify whether a presented object was concave or convex (see [3] for a complete description of this experiment and many other analyses of both the behavior and the neural activity). Each trial began with an object (convex or concave) moving toward the whiskers (*t* = −2 sec). Objects could stop at one of three different distances (far, medium or close), which happened at *t* = −0.9, *t* = −0.7 and *t* = −0.5 seconds, respectively. The uncertainty about distance made the task more difficult, and it encouraged the animals to use all the whiskers at their disposal (C0, C1, C2, and C3). When the response window opened (*t* = 0 seconds), mice had to make a choice by licking the left lickpipe for concave objects and the right lickpipe for convex objects. The object position and the whiskers were monitored using high-speed video and processed with a deep neural network [20, 21, 22] (Fig. 2b; see Methods). Importantly, mice were free to whisk and lick throughout the course of the trial (2 seconds). On each trial, the choice of the animal was determined by the side of the first lick after the response window opened at *t* = 0 seconds.

**Figure 2:**
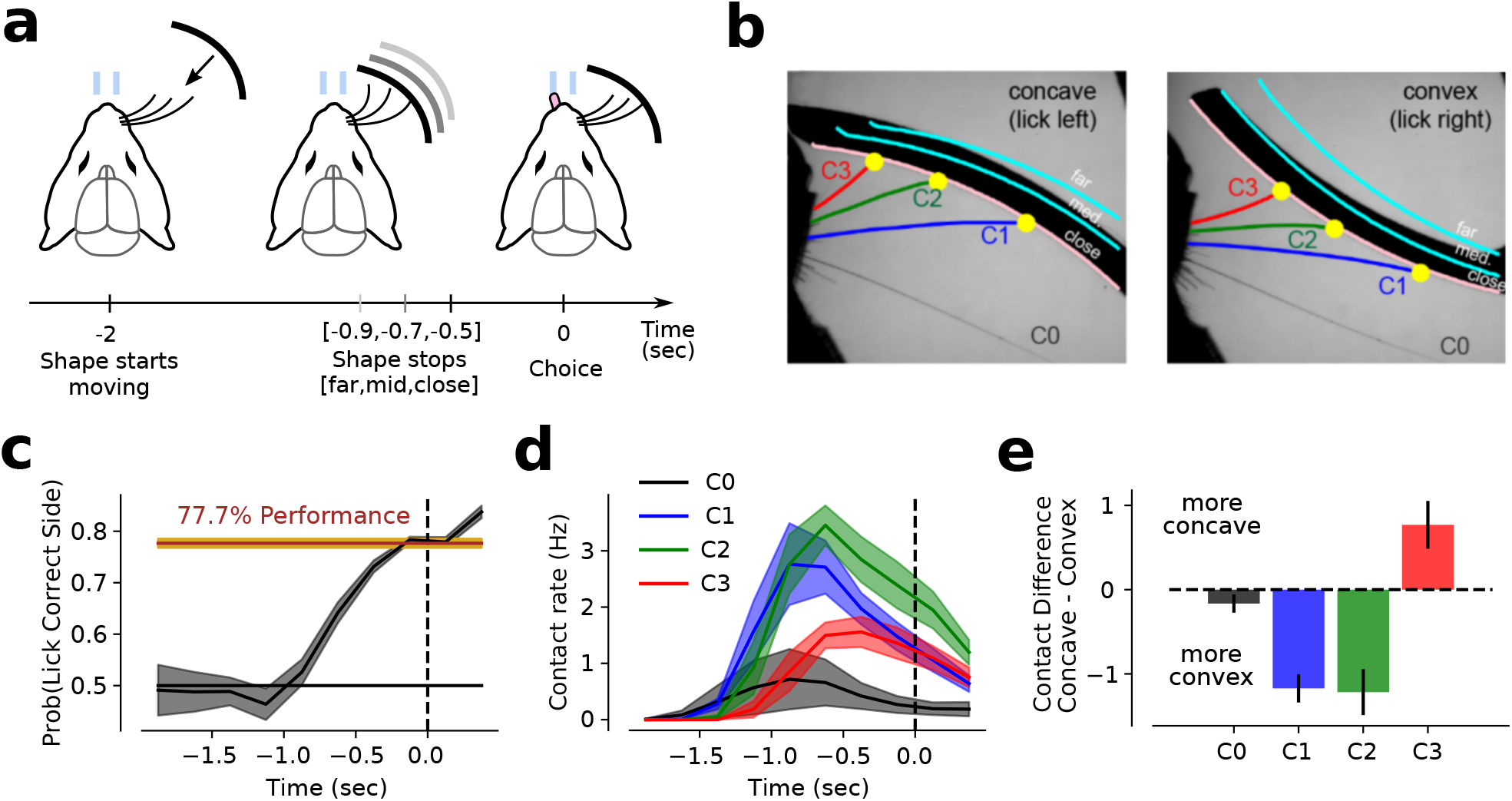
Mice gather whisker evidence throughout the trial to discriminate concave from convex shapes. (**a**) Animals were presented with either a convex or a concave shape and after two seconds they had to report their choice by licking the left (concave) or right (convex) lickpipe. (**b**) Whiskers and shape position were monitored by combining a high-speed camera with an image parsing algorithm [20, 21, 22]. (**c**) The probability of making a lick on the correct side (y-axis) increased as a function of time (x-axis) throughout the trial. Animals had a mean performance of ~ 78%, defined as the probability of making a correct lick at the moment of choice. (**d**) The time profile of contact rates increased significantly once the shape was within whisking distance and it was similar across all whiskers (C0, C1, C2 and C3). (**e**) Difference in the total number of contacts between concave and convex shapes (y-axis) for all whiskers. C1 and C2 made more contacts for convex shapes, while C3 made more contacts for concave shapes. Panels (c-e) were obtained using a sliding window of 250 milliseconds. Errorbars in (c-e) correspond to s.e.m. across animals.

Mice performed the task with a mean accuracy of 77.7% ± 0.9% (s.e.m.) (Fig. 2c). The probability of making a lick on the correct lickpipe increased throughout the course of the trial, indicating that mice based their decision on the accumulated sensory evidence gathered by whisking. This implies some form of temporal integration. Most contacts were made between *t* = −1.25 and *t* = −0.25, suggesting this was the most informative time window (Fig. 2d).

The whiskers contacted the two shapes at different rates: the mean difference in total number of contacts between convex and concave objects for whisker C1 was −1.17± 0.17, for C2 −1.22 ± 0.27 and for C3 0.77 ± 0.28 (Fig. 2e). Therefore, by computing the weighted sum of total number of contacts of the three whiskers, it should be possible to discriminate between convex and concave objects. The contact rate of each whisker followed a similar time profile (Supplementary Fig. S1). Mice made more contacts on correct trials (Supplementary Fig. S2), suggesting that errors resulted from poorer sensory gathering or a lower level of task engagement.

### Linear integration is sufficient for object discrimination

In order to understand whether linear integration is actually sufficient to determine the curvature of an object, we first predicted stimulus shape on a trial-by-trial basis using a linear decoder that reads out the spatio-temporal pattern of whisker contacts observed during the execution of the task (Fig. 3a). As whisker contacts were not necessarily the only behaviorally-relevant variables, we also included other variables like the angular position of the whiskers during contacts. Moreover, we were interested in understanding whether these spatio-temporal patterns could explain the decision of the animal, which not always matches the shape of the object (the animals make mistakes). So we also trained a decoder to predict the side where the animal licked at the response time (choice). Both predictions were tested on held-out trials (cross-validation; see Methods).

**Figure 3:**
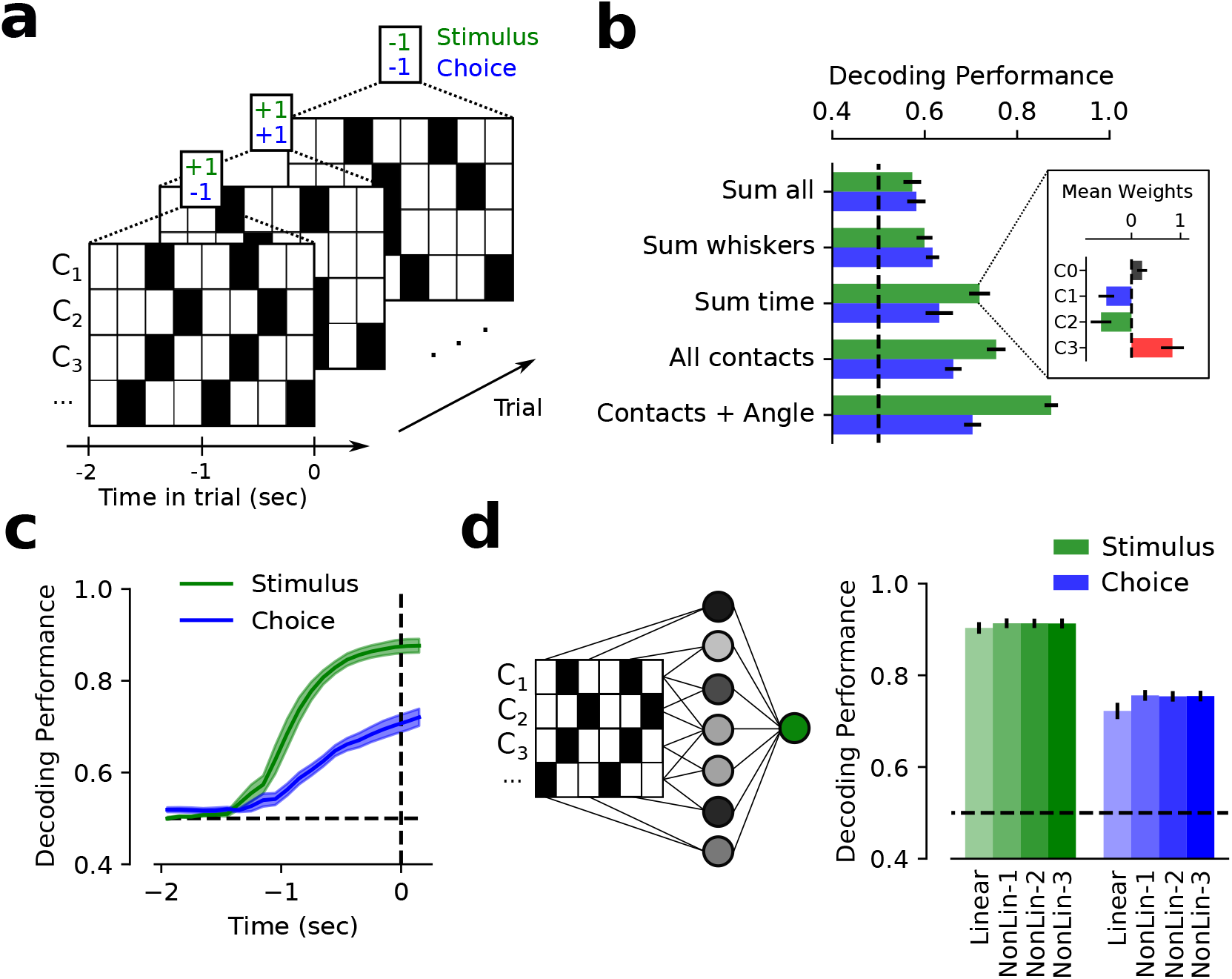
The whisker-based shape discrimination task can be solved by linearly integrating whisker contacts across time. (**a**) The spatio-temporal recorded pattern of contacts across time and whiskers was used to classify shape identity (green) and animal choice (blue) on a trial-by-trial basis. (**b**) Stimulus and choice decoding performance when different input features were used. Sum all: the decoder used only the sum of whisker contacts across all time bins and whiskers; Sum whiskers: sum across whiskers, but the temporal structure of the sum is retained; Sum time: the decoder considers the vector of the total number of contacts for each whisker; All contacts: the decoder reads out the full spatio-temporal pattern of all contacts from all time bins and whiskers; Contacts+Angle: all contacts and the angle of the whisker at time of contacts, which produces the highest decoding accuracy. (Inset) The weights obtained by a classifier trained to decode shape identity match the difference in number of contacts between concave and convex shapes (see Fig. 2e). (**c**) Decoding performance as a function of time when the decoder reads out the full spatio-temporal pattern of contacts and angle from −2 seconds to the time indicated on the x-axis. The performance increases as a function of time for both shape identity and animal choice. (**d**) A multi-layer neural network model is trained to use the full spatio-temporal pattern of contacts and angle of contact to predict the stimulus and the choice of the animal. Non-linear, multi-layer neural networks perform similarly to the linear network with no intermediate layer. Thus, linear integration of the sensory cues is sufficient to predict both stimulus and choice. Error bars in all panels correspond to s.e.m. across mice.

The most informative set of features comprised all the whisker contacts and angle of contact across time (Fig. 3b, see also [3]). When whisker contacts were summed either over time, or across whiskers, the performance was lower, indicating that the full spatio-temporal pattern of whisker contacts best predicts the curvature of the stimulus and the choice of the animal. Unsurprisingly, the weights of the classifier trained on contacts summed over time (Fig. 3b inset) reflected the difference in total number of contacts for convex vs concave objects (Fig. 2e). We also observed that the accuracy of the classifiers increased as mice accumulated more evidence (Fig. 3c; see Methods). This observation nicely matches the progressively increasing probability of licking on the correct side of Fig. 2c.

Until now we only considered simple linear decoders, which compute a weighted sum of the inputs (whisker contacts and angles of contact at different times) and compare it to a threshold. We next asked whether the shape discrimination task could be better performed by non-linearly integrating some of the features that characterize the sensory input. We also asked whether the observed behavior was better explained by a linear or a non-linear sensory integration process. In other words, we assessed whether linear or non-linear decoders would better predict the stimulus identity and animal choice on a trial-by-trial basis. We trained two different feed-forward neural networks with non-linear units arranged in multiple layers to predict stimulus and choice, respectively. These decoders are more complex and they contain more parameters than the linear ones, so they will certainly perform better at classifying the patterns in the training set. However, it is not guaranteed that the cross-validated performance, computed on held-out trials, will actually increase. The cross-validated performance of the non-linear classifier can only surpass the linear classifier if non-linear combinations of the features are important, indicating that the task could be solved more efficiently by combining the inputs variables in a non-linear way.

Despite the task being significantly more complex than others considered in the past, we observed that both linear and non-linear decoders performed similarly at classifying stimuli (linear: 90.3% ± 1.3%; best non-linear: 91.3% ± 1.1%) (Fig. 3d; green). A similar result was observed on both correct and error trials (Supplementary Fig. S3a), though for error trials the performance was significantly lower for all decoders. This performance decrease is likely due to the lower number of contacts and overall lower task engagement in error trials (Supplementary Fig. S2). When predicting choice on a trial-by-trial basis, a similar trend was observed (linear: 72.2% ± 1.8%; best non-linear: 75.6% ± 1.2%) (Fig. 3d; blue), suggesting that animals’ decisions were mostly driven by a linear combination of the sensory cues across time and whiskers. On trials when the mouse made a mistake, our ability to predict its choice was substantially lower, suggesting that these trials were qualitatively different (Supplementary Fig. S3a). It is important to note that all models were trained on a balanced dataset in which the number of correct and error trials was exactly the same. This procedure decorrelates the variables stimulus and choice, which otherwise would be highly correlated. Finally, in order to discard the possibility that the results in Fig. 3d were a consequence of a low number of trials, linear and non-linear decoders were also trained on synthetic tasks of whisker contacts with different levels of difficulty (see Methods). Unsurprisingly, linear and non-linear decoders performed equally well on simple integration tasks, whereas non-linear decoders were necessary to perform complex tasks that require non-linear integration of sensory evidence (Supplementary Fig. S3b).

### Non-linear mixed selectivity in the mouse S1 cortex

To characterize how task variables are represented in the somatosensory cortex (S1) of behaving mice, multiple neurons were simultaneously recorded in this brain region while mice performed the whisker-based object discrimination task (Fig. 4a). S1 neurons were predictive of shape identity and animal choice on a trial-by-trial basis as revealed by the performance of a linear classifier (Fig. 4b left panel, see Methods). At response time, shape category could be decoded from small ensembles of neurons (mean population size of 25.4 cells) with a performance of 56.8% ± 1.7% (green) and the choice of the animal with a performance of 65.4% ± 2.0% (blue). Both shape and choice could be reliably decoded from neuronal ensembles in which we grouped together the activity from different recording sessions (pseudopopulations)[3]. As expected, populations of S1 neurons encoded information about whisker contacts (Fig. 4b right panel), where the task was to predict whether a particular activity pattern corresponded to a trial with high or low number of contacts for each whisker (see Methods).

**Figure 4:**
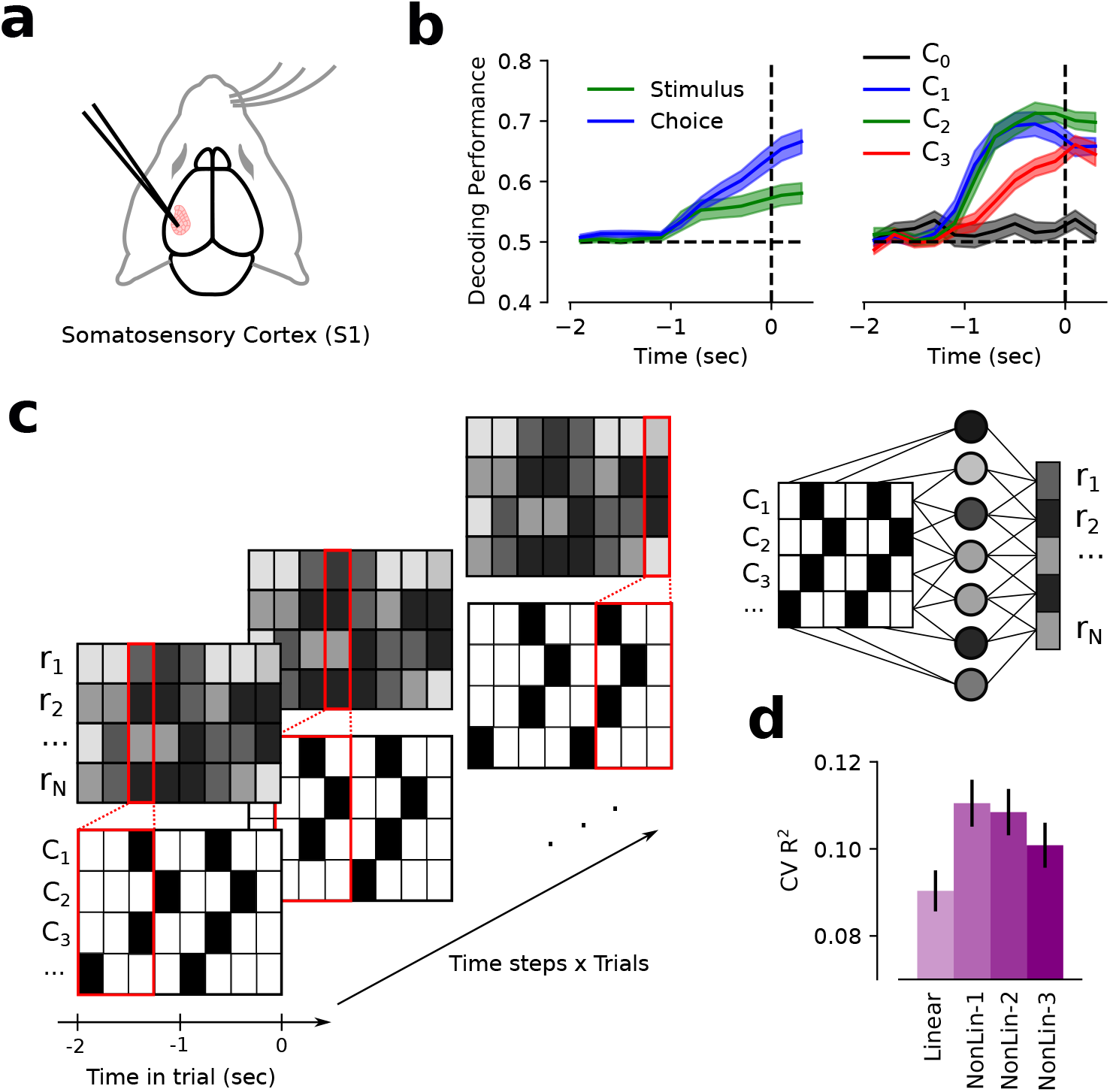
Populations of neurons in the mouse somatosensory cortex (S1) exhibit nonlinear mixed selectivity for task variables. (**a**) Populations of S1 neurons were simultaneously recorded while mice performed a whisker-based discrimination task. (**b**) Decoding performance of a linear classifier (y-axis) that reads out from S1 to predict stimulus or choice (left panel), or contacts made by each whisker (right panel) as a function of time (x-axis). For each timepoint, decoding performance was evaluated using all neuronal activity time bins from −2 seconds to the time indicated on the x-axis. As expected, decoding performance is at chance early in the trial when mice do not make contacts and it reaches ~65% at the end of the trial for choice (blue), and ~57% for stimulus (green). Whether whiskers made high or low number of contacts on a given trial could be decoded more accurately than shape identity. (**c**) S1 activity was regressed against task variables. Linear and nonlinear neural network models with a different number of intermediate layers were used to reproduce the observed neural activity. (**d**) Cross-validated *R*^2^ (y-axis; see Methods) on populations of S1 neurons for the different encoding models (x-axis). A fully connected neural network with one hidden layer (NonLin-1) outperforms the linear model and other neural networks with more intermediate layers (left panel). Errorbars correspond to s.e.m. across neurons.

To determine whether the neurons responded also to other variables, we trained encoding models that predicted the firing rate of the recorded population from the set of whisking and behavioral variables for all time steps (100 ms) and trials (Fig. 4c). In particular, we used as regressors of the encoding models instantaneous task variables like whisking contacts and angle, lick side and rate, as well as trial task variables like current and previous reward, choice and stimulus (see Methods). Fitting encoding models is useful when it is unclear which variables are modulating the activity of a population of neurons. Moreover, a mapping between task variables and neural activity can be understood as a multi-dimensional generalization of the tuning curve of a population of neurons and it is also useful for constructing denoised (smoothed) neural representations.

We considered four different encoding models that were implemented using feed-forward neural networks: one that simply implements linear regression (0 hidden layers), and three feed-forward neural networks with 1, 2 and 3 hidden layers of rectified linear units (ReLU) and linear output. Note that the linear regression network can only generate pure and linear mixed selectivity neurons.

In contrast, the 1, 2, and 3-layer neural nets can generate neuronal responses that depend on nonlinear interactions between the different regressors (non-linear mixed selectivity). We found that the activity of populations of S1 neurons was best explained by a non-linear mixed selectivity encoding model (1-layer neural network; *R*^2^ = 0.111 ± 0.005 on held-out data; see Methods) (Fig. 4d). The linear model with only pure and linear mixed selectivity was the worst at explaining the neural data (*R*^2^ = 0.089±0.005). Including intermediate layers in the encoding model produced an increase of 24% in explanatory power. As expected, when all models were tested on training data, more parameters entailed better firing rate prediction (Supplementary Fig. S4a). Models evaluated on correct trials showed better performance than those evaluated on incorrect trials, likely due to the reduced number of trials and the smaller number of contacts made on mistakes (Supplementary Fig. S4b). The encoding models showed a higher performance for inhibitory neurons and neurons located in deeper layers of the somatosensory cortex (Supplementary Fig. S5), possibly due to their higher firing rates. All of these results were qualitatively equivalent when a Poisson loss function was used instead (Supplementary Fig. S6).

To assess the importance of each regressor, we calculated 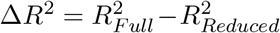, which represents the loss in prediction power on held-out data when a particular regressor or group of regressors is set to zero (Fig. 5a, see Methods). Whisker contacts and continuous whisker angular position were the two most important variables for explaining the neuronal responses (Fig. 5b). Interestingly, superficial layers (2/3) were more strongly driven by sensory (contacts) than motion variables (whisker position), while deep layers (5 and 6) showed the opposite trend (Supplementary Fig. S7). As expected, the time kernel for whisker features showed a recency effect for all whiskers. Population activity was better predicted by C1 and C2 contacts than by C3 contacts, and it was less well predicted by the angular position of C1 than by the other whiskers (Fig. 5c). In agreement with [3], we also found a deviation from classic somatotopy: C1 contacts were more strongly represented than C2 contacts in the C2 column, and than C3 contacts in the C3 column (Supplementary Fig. S8). Previous reward *R*_−1_ had the strongest effect before whisker contacts were typically made (−2 sec to −1 sec), while the current reward *R*_0_ peaked after the response window opened and the animal made its choice (Fig. 5d). Although the current stimulus *S*_0_ (shape category) and choice *C*_0_ (lick side) followed a similar trend throughout the course of the trial, *C*_0_ had a stronger effect on the neural activity at the beginning of the response window. Similar task variable time profiles have been reported in previous studies in other animals and brain regions [7].

**Figure 5:**
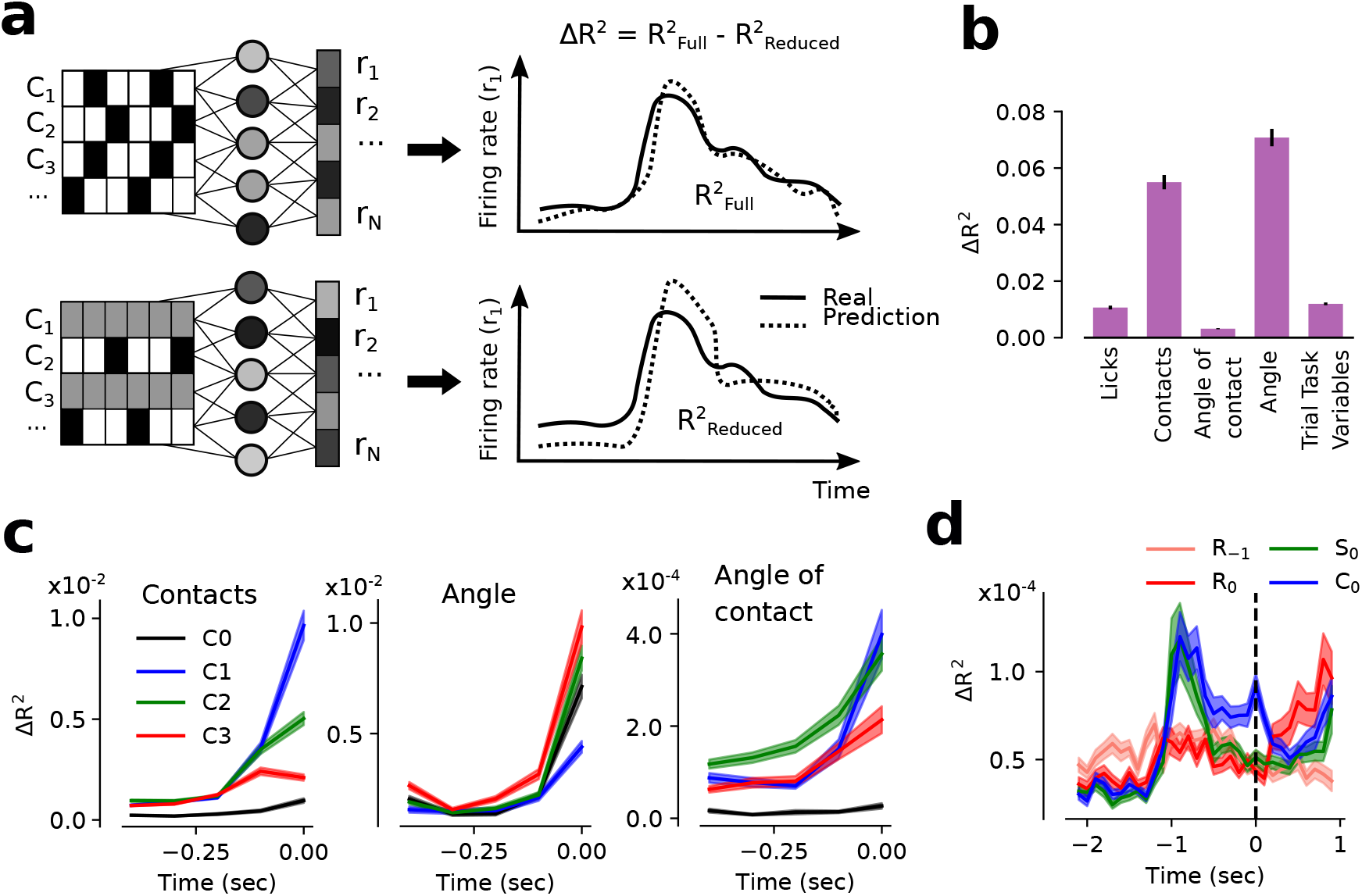
Contacts and whisker angular position are the variables that contribute the most to the prediction of S1 population activity. (**a**) We fit an encoder model to explain the population’s firing rate (*r*_1_, *r*_2_,…, *r_N_*) as a non-linear function of task variables like whisker C1, C2 and C3 contacts (i.e. *C*_1_, *C*_2_ and *C*_3_), and calculated 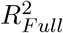, the goodness-of-fit of the full model (top). To assess the importance of each regressor, we set the input data for that regressor (or group of regressors) to zero (gray) and assessed the goodness-of-fit of the reduced model, 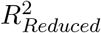 (bottom). In this way we quantified the importance of each regressor as the resulting decrease in goodness-of-fit 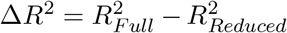. (**b**) Whisker contacts and angular position were the most important factors on S1 activity as revealed by the decrease in model accuracy Δ*R*^2^. (**c**) Decrease in model accuracy (y-axis; Δ*R*^2^) for whisker contacts (left panel), angle of contact (central panel) and angular position (right panel) for different time lags with respect to current time step (x-axis). Neural activity was better explained by variables in the current time step. (**d**) Previous reward *R*_−1_ was predictive of neuronal activity (y-axis; Δ*R*^2^) early during the trial whereas the importance of current reward *R*_0_ peaked after mice made their choice. Additionally, although current stimulus *S*_0_ and choice *C*_0_ followed a similar trend throughout the course of the trial, *C*_0_ had a stronger effect on the population’s firing at response time (t = 0). Errorbars in all panels correspond to s.e.m. across neurons.

### What does somatosensory cortex mix?

The recorded neurons displayed a wide range of response properties: while some neurons showed approximate linear mixed selectivity for C1, C2 and C3 contacts (Fig. 6a), others showed sub-linear or XOR-like responses (Fig. 6b). To study non-linear mixed selectivity we could have fit a linear model with interaction terms for each neuron. Instead, we decided to fit the neural network encoding model to all the neurons simultaneously recorded, so that we could determine in an unbiased way whether and which interaction terms were shared by multiple neurons. The state of activation of each unit in the intermediate layer is a weighted sum of the inputs passed through a non-linearity. Thanks to the non-linearity, the units’ responses contain interaction terms. During training, the weights of the intermediate units are tuned to produce the interaction terms needed to explain the neuronal responses. These terms are then available to all the neurons whose activity we intend to reproduce.

**Figure 6:**
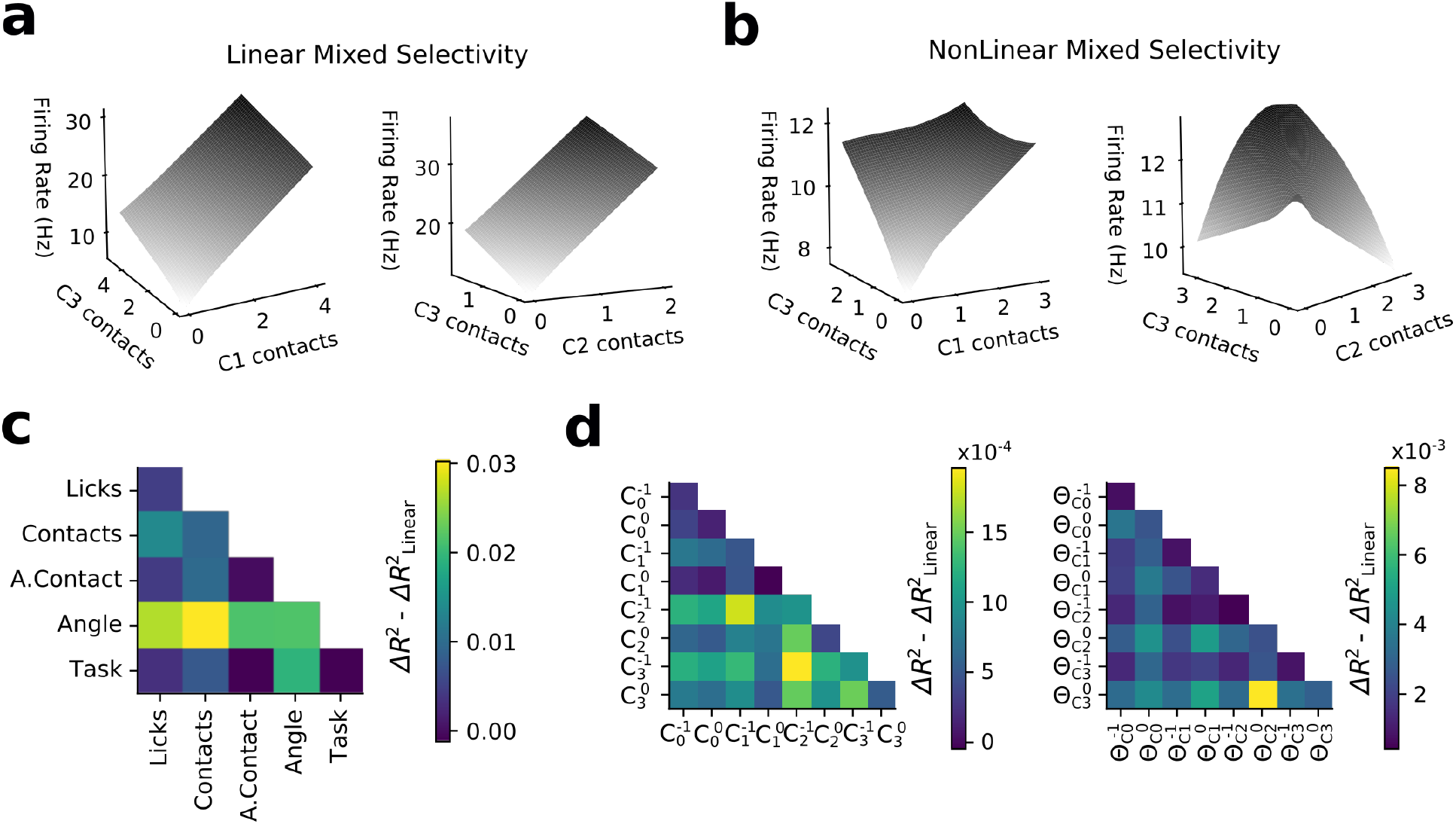
Neurons show non-linear mixed selectivity, mostly originating from non-linear interactions between contacts and whisker angular position. (**a-b**) Tuning curves of example neurons that exhibit approximately linear mixed selectivity (a) and non-linear mixed selectivity (b) for C1, C2 and C3 whisker contacts. All tuning curves were obtained from the best encoding model (non-linear with one hidden layer, see Fig. 4d). (**c**) Non-linear mixed selectivity 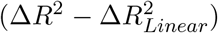 for the interaction between the different groups of task variable. The interaction between whisker contacts and angular position was the most important non-linear contribution to the encoding model. Diagonal elements account for the strength of the non-linearity between a particular task variable and the activity of the neuronal population. (**d**) Non-linear mixed selectivity contribution for different time steps and whisker contacts (left) and angular position (right). The strongest interaction for contacts between whiskers occurs at time lags of 100ms (e.g. 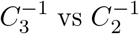), while for angular position it occurs at time lags of 0ms (e.g. 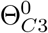 vs 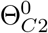) between neuronal activity and task variables.

Once the network was trained, we determined the contribution of non-linear interactions of specific pairs of variables by setting the two variables to zero and evaluating both Δ*R*^2^, which is the loss in explanatory power for the full non-linear model (see Fig. 5b-d) and 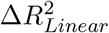, which is the analogous loss for the linear model. We then computed the difference 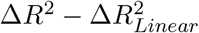 (see Methods and Supplementary Fig. S9). If this difference is close to zero, then the interaction term under consideration is not important, because the loss is the same whether we use the full non-linear model, which can compute the interaction term, or the linear model, which cannot. If the difference is large, then the interaction term is important.

We found that the most important interaction was between whisker contacts and angular position (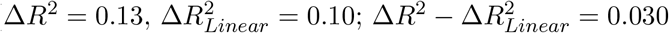; Wilcoxon signed-rank test, *P* < 0.001; Fig. 6c), followed by interactions between whisker angular position and the rest of the variables. Similar results were observed in both excitatory and inhibitory neurons, and across layers (Supplementary Fig. S10). We next assessed the importance of the interactions between different whiskers and time steps (time lags) for the task variables whisker contacts and angular position. For contacts, the interactions were strongest not for the current time step, but when the preceding time bin of 100ms was considered. This is an interesting and non-trivial result given that contacts at the current time steps are those that most affect the neural activity (Fig. 6c) and could be reflecting the effect of response inhibition between whisker contacts that occur within whisk cycles of 50-100 ms. For the angular position of the whiskers, the strongest interactions were observed in the current time bin (Fig. 6d).

We also examined non-linear interactions between contacts and whisker position across all whiskers and at multiple time steps. Interestingly, the strongest non-linear mixing between angular position and contacts also occurred with a time lag of 100ms (Supplementary Fig. S11; see Discussion for the implications of these observations). Finally, similar to how neurons in C2 and C3 columns showed an unexpected response to C1 contacts (Supplementary Fig. S8), we observed that they also responded to non-linear interactions with the C1 whisker, regardless of their anatomical location (Supplementary Fig. S12).

### The geometry of the neural representations in S1

Representations generated by models that include non-linear interactions between task variables like whisker contacts and angular position turned out to be the best description of the activity in S1. However, this finding describes only one aspect of the geometry of the representations in S1. Indeed, almost any geometry that is different from the low-dimensional geometry G1 (Fig. 7a) would be compatible with the observed non-linear responses of individual neurons. To better characterize the observed representational geometry in S1, we studied how it could be used by a downstream linear readout, which could be interpreted as a downstream neuron. Specifically, we trained the linear readout on the S1 activity to perform different synthetic tasks, each requiring different properties of the geometry of the representation. For instance, when the representations are low-dimensional (G1; Fig. 7a) a linear readout can perform well on easy tasks. In these situations the readout can also easily generalize to novel situations. However, low-dimensional representations lead to poor performance in complex tasks, which require the ability to discriminate between non-linear combinations of the different task variables. On the other hand, high-dimensional geometries (G3) are well suited for complex tasks but their ability to generalize is limited. Moreover, the fact that noise is typically amplified in high-dimensional representations, can impact negatively the performance on simple tasks. We propose that a geometry in between G1 and G3 could enable a linear readout to discriminate without affecting much the ability to generalize (G2).

**Figure 7:**
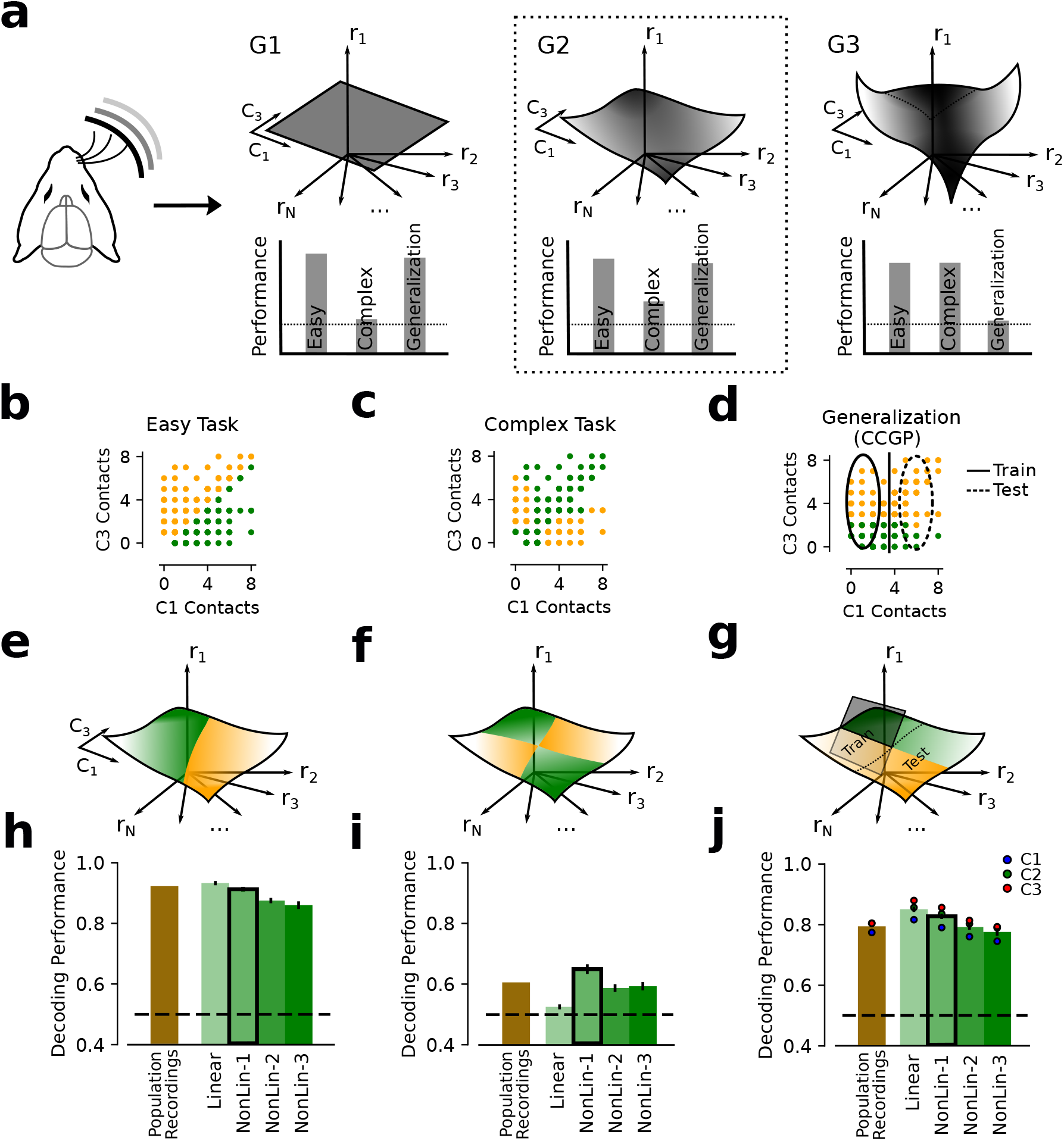
The representational geometry in S1. (**a**) A linear readout performs and generalizes differently on neural representations with each of the three geometries introduced in Fig. 1. For low-dimensional representations (G1) in the neural activity space, *C*_1_ and *C*_3_ are represented along orthogonal axes. For these representations, the performance is high in easy tasks and the readout can readily generalize to novel situations. However, the performance is poor in complex tasks that require non-linear combinations of *C*_1_ and *C*_3_. High-dimensional representations (G3) allow for high performance in complex tasks, but generalization is poor. Intermediate geometries (G2) could benefit from the computational properties of both low- and high-dimensional representations. (**b**) Easy task: the linear classifier has to output 1 (orange) if the weighted sum of *C*_1_ and *C*_3_ is larger than a threshold, and 0 otherwise (green). The weights were random and the two classes are linearly separable. The values of *C*_1_ and *C*_3_ (or other pairs of whisker contacts) are taken from the experiment. (**c**) Complex tasks: the (*C*_1_,*C*_3_) space is divided into 4 regions by two orthogonal random directions. The two classes, each comprising two diagonally opposite regions, are colored in orange and green. The task is not linearly separable. (**d**) Generalization benchmark: the cross-condition generalization performance is measured for both *C*_1_ and *C*_3_. In the figure we show the CCGP for *C*_3_: a linear decoder is trained to discriminate between high and low *C*_3_ on the data points on the left (Train; low *C*_1_), and it is tested on the data points on the right (Test; high *C*_1_). (**e**-**g**) The geometry G2 for the different tasks. For each combination of *C*_1_ and *C*_3_ we used the real neural data (pseudosimultaneous recordings) or the data generated by linear and non-linear encoding models. (**h**) Easy task performance: high with real data (brown bar). The performance is highest for the linear encoder, though the non-linear encoders perform almost as well. (**i**) Complex task performance: above chance, except for the linear encoder representations. (**j**) Generalization: CCGP is high for all whisker contact variables (dots; different axes in g) and for all the surrogate data generated by the different encoders. Errorbars in all panels correspond to s.e.m. across populations of simultaneously recorded neurons (recording sessions).

To study the geometry of the neuronal representations in S1, we started from the real spatiotemporal patterns of whisker contacts and we defined a series of synthetic classification tasks. More specifically, the desired output of the linear readout was decided on the basis of the vectors containing the total number of whisker contacts in a trial, for three whiskers. We then considered the recorded neural representations constructed by combining together all the neurons from different sessions (same or different animals) and concatenating the activity vectors of all time bins within a trial (pseudopopulations; see Methods), and we trained a linear classifier to perform the classification task by reading out these neural populations. To better understand the contribution of linear and non-linear components of the activity, we also performed the same analysis on the neural representations generated by the different encoding models described in the previous section.

We considered complex and easy tasks, and for an easy task we estimated the ability to generalize in novel situations. Both the complex and the easy tasks are binary classifications tasks of the number of whisker contacts. To define the two classes we considered two whiskers at a time and we ignored the number of contacts of the third whisker. For the easy task we computed the weighted sum of the number of contacts for two whiskers (e.g. C1 and C3, as in Figure 7b). The weights were random, and each choice of weights corresponded to a different implementation of an easy task. The class to which a particular input (pair of C1 and C3 contacts) belongs depended on whether the weighted sum was above or below a threshold. In Figure 7b we indicated one class with green and the other with orange. The two classes are separated by a line (not shown) whose orientation depends on the random weights. By construction, the easy task is linearly separable. For the complex task, the space of whisker contacts was divided into 4 regions by two orthogonal separating lines, again in random directions. Each class included the points of two diagonally opposed regions (Fig. 7c). This task is similar to a XOR task, which is known to be non-linearly separable. Finally, to benchmark the ability to generalize, we computed what is called cross-condition generalization performance (CCGP), which is a signature of a process of abstraction [4]. CCGP was evaluated as a linear decoder’s ability to report the number of contacts (high vs low) of one whisker from the neural activity, when trained on a subset of cases (see Methods). For example the decoder is trained to report C3 contacts when the number of C1 contacts is small, and then it is tested in the cases in which the number of C1 contacts is large (Fig. 7d). If C1 and C3 contacts are represented in approximately orthogonal subspaces, then the CCGP is high. This geometry allows generalization to novel situations (for example, the high C1 contact case is novel for the decoder trained to report C3 contacts only when the number of C1 contacts is small).

For the pseudopopulation recordings, we observed that the linear decoder could perform the easy task with high accuracy (brown bar; Fig. 7h). The performance for the complex task was reduced, but still above chance (brown bar; Fig. 7i). Interestingly, CCGP was high for all the variables representing the number of contacts of different whiskers (brown bar; Fig. 7j). This implies that the number of contacts for different whiskers are represented in approximately orthogonal subspaces of the neural activity space. Moreover, it means that the coding direction for the number of contacts of each whisker does not depend much on the number of contacts of the other whiskers. In other words, the coding directions of each whisker number of contacts are parallel to each other when one considers different values of the number of contacts of the other whiskers. Importantly, we found that all columns in S1 encoded information about all whiskers in approximately orthogonal subspaces (Supplementary Fig. S13).

To better understand the role of non-linearities in the responses of individual neurons, we then computed the performance of the linear decoder on easy and complex tasks and we estimated the CCGP when the neural representations are not those recorded, but the representations generated by linear and non-linear encoders (Fig. 7e-g). Given that all the encoding models were fit using time windows of 500ms, the easy and the complex task in this case were defined with respect to the integrated number of whisker contacts in time windows of 500ms. As expected, for the easy task, the linear representations worked slightly better than non-linear representations (green bars; Fig. 7h). Non-linearities can lead to noise amplification, which can impair performance on simpler tasks. For the linear encoding representations, the linear decoder could not perform the complex task. The representations generated with the non-linear encoding model with only one hidden layer produced the best performance on the complex task (green bars; Fig. 7i). Finally, not surprisingly, CCGP was higher for linear representations, which are lower-dimensional. However, once again, as in the case of easy tasks, the advantage was modest (green bars; Fig. 7j). Performance on the easy and complex tasks as well as CCGP depend on the number of neurons (Supplementary Fig. S14). Larger populations of neurons produce a higher decoding performance, which is particularly relevant for the complex task. Qualitatively equivalent results were also obtained when the synthetic tasks were defined with respect to continuous whisker angles or the interaction between whisker contacts and continuous angles (Supplementary Fig. S15).

Overall, these results indicate that the geometry of the representations in S1 is best described by G2 in Figure 7a, which is obtained by starting from a low-dimensional scaffold (e.g. the square of G1) and perturbing it with non-linearities. The non-linearities enable a linear readout to perform complex tasks and the low-dimensional scaffold allows for high generalization performance, as measured by CCGP. The non-linearities reduce only slightly the robustness to noise in easy tasks, but they significantly increase the ability to perform complex tasks.

### Task difficulty modulates the geometry of the representations in RNNs

In the previous section we have seen that the neuronal representations in mouse S1 contain a lowdimensional scaffold, which allows for generalization, but also significant non-linearities that enable a linear readout to perform complex tasks (G2 in Fig. 7a). In Fig. 3 we also showed that the whisker-based discrimination task can actually be solved by a linear combination of whisker contacts, which implies that a low-dimensional geometry (G1) in S1 would be sufficient to perform this task. If a low-dimensional geometry suffices, then why would neuronal representations in S1 include nonlinearities between task variables (G2)? The same neural representations are probably employed in a number of different behaviors beyond the one studied in the experiment. Thus, we next asked how the geometrical properties of the representations are modulated by the task needs.

We trained recurrent neural networks (RNNs) to perform tasks with different levels of difficulty that are similar to the shape discrimination task (Fig. 8a). The reason we studied RNNs is that they can integrate information over time, as required by these tasks and by the experimental task. Importantly, the RNNs were only required to generate the correct response in the artificial tasks, and not to reproduce the neural data [23, 24]. The different tasks had the following structure: each trial lasted 30 time steps, and at each time step we fed into the RNN a vector containing three binary variables (each representing contacts made by one of three whiskers). Each binary variable was random (an independent Bernoulli process) with either a high or a low success rate λ, for a total of 8 different input patterns. The desired outputs defined three tasks with different levels of difficulty: easy (Fig. 8b), medium (Fig. 8c) and complex (Fig. 8d). The easy task could be solved by linearly mixing information across input channels whereas the medium and complex tasks required non-linear mixing of sensory cues by the artificial units. We trained a different network to perform each of the three tasks, and for each network we trained input, recurrent, and readout weights.

**Figure 8:**
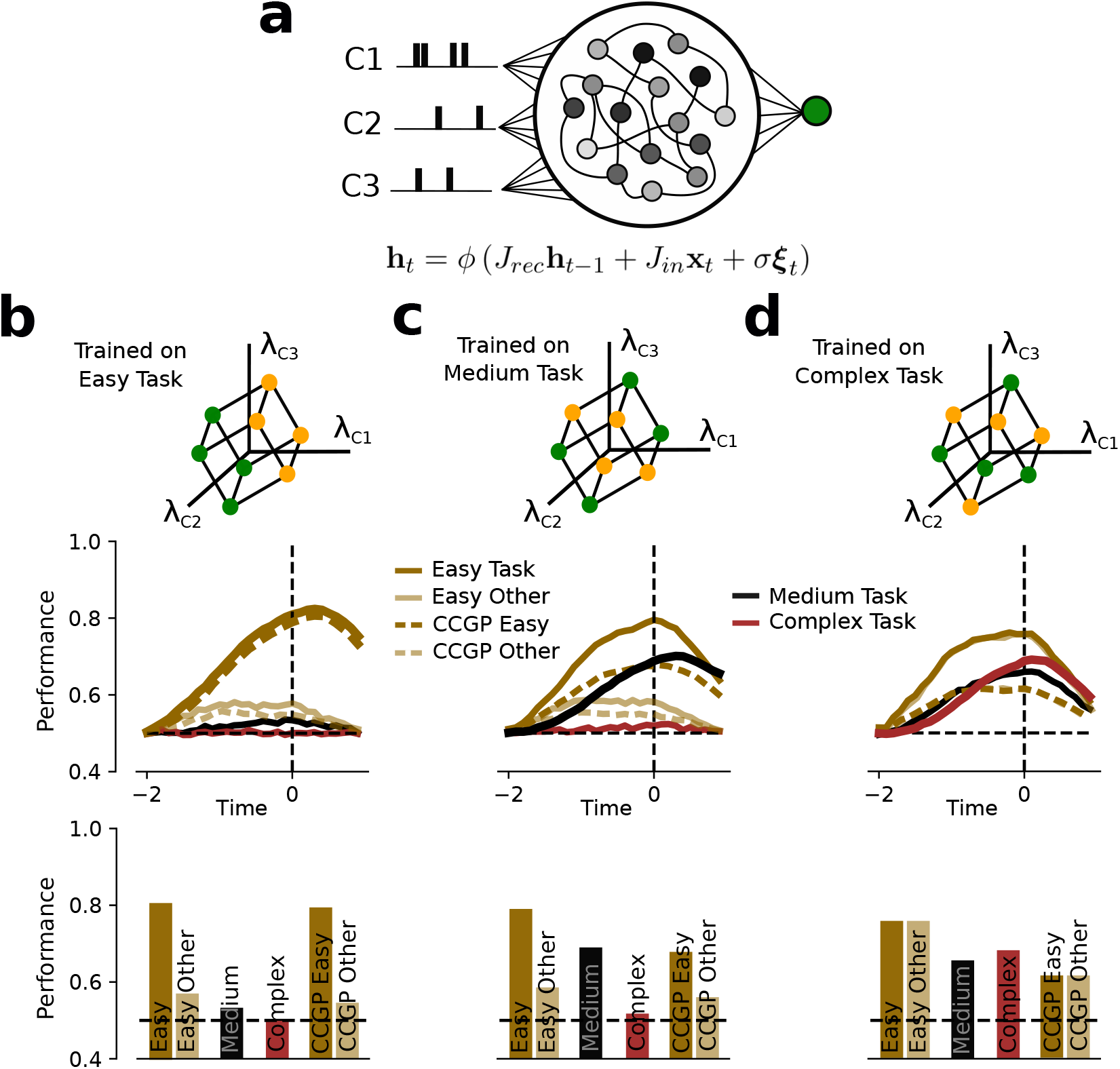
The geometry of the representations in RNNs is modulated by the difficulty of the task. (**a**) A set of noisy and fully connected ReLU units receive input from three independent channels (C1, C2 and C3). On each trial, each channel corresponded to a Bernoulli process drawn from either a low or a high success rate parameter λ. The state of the network at a given time step (**h**_*t*_) is determined by its state at the previous time step (**h**_*t*−1_), the input at the current time step (**x**_*t*_) and independent Gaussian noise (**ξ**_*t*_) (see Methods). (**b**) Top panel: the easy task consists of a linear integration task across input channels. Middle panel: probability correct (y-axis) as a function of time within a trial (x-axis). Due to the inherent stochasticity of the input trains, performance increased as a function of elapsed time in all networks and tasks. When the network is trained on the easy task (dark brown), a readout unit fails at performing an orthogonal easy (easy other; pale brown), a medium (black) and the complex task (maroon) (see Methods). Generalization, defined as the cross-condition generalization performance (CCGP), is high for the easy task and low for the orthogonal other easy task. Bottom: profile of performances of the “Easy Task” RNN at choice time (*T* = 0) for the different discrimination and generalization tasks. (**c**) Top panel: the medium task consists of a non-linear integration task with respect to two input channels (2D-XOR). Middle panel: when the network is trained on the medium task, the representation allows a readout unit to also perform the easy task but fails at the orthogonal other easy and complex tasks. Generalization for the easy task is higher than for the easy orthogonal task. Bottom: same as (b) for the “Medium Task”. (**d**) Top panel: the complex task consists of a non-linear integration task across all input channels (3D-parity). Middle panel: training a network on the complex task produced a high-dimensional representation that allowed a readout unit to perform all the easy and the medium tasks. The representation of the variables that defined the easy tasks allowed for generalization well above chance (CCGP). Bottom: same as (b-c) for the ‘‘Complex Task”. Different levels of input noise produced qualitatively equivalent results for all networks, tasks and generalization (Supplementary Figs. S16 and S17). For all panels, the performance curves correspond to the mean across 50 random realizations of input patterns and tasks (see Methods).

To study the geometry of the neural representations of the different RNNs, we adopted an approach similar to the one we used for real data: we trained a linear readout to perform different tasks. We asked how well the representations learned during training on one task could be used to perform other tasks. Specifically, we trained each network on a single task (easy, medium, or complex), froze the recurrent and input weights, and then optimized linear readouts to use those representations to perform each of the other tasks. Moreover, for the easy and medium tasks, we considered all possible equivalent tasks that correspond to the separation of different groups of points (easy other and medium other; Fig. 8 and Supplementary Fig. S16). For example, there are three different easy tasks, each separating the 8 points into the two groups that correspond to opposite faces of the cube (see also Methods). To study the generalization properties of the neural representations we estimated the CCGP for different dichotomies (ways of dividing the points into two groups) that correspond to different easy tasks (Fig. 8 and Supplementary Fig. S17).

Networks trained on an easy task produced high discrimination and generalization performances (high CCGP) for the easy task only (dark brown; Fig. 8b), suggesting that the network had learned a low-dimensional representational geometry that was specialized for this simple task. Training a network on the complex task produced representations that allowed a linear readout to perform all the different tasks (Fig. 8d) and interestingly the easy tasks could be performed better than the task the network was trained on. Importantly, CCGP was well above chance for all the variables defined by the three possible easy tasks. When the RNN was trained to perform the medium task (Fig. 8c), the representational geometry was well suited to discriminate and generalize for the easy task. Of course it also performed well the task it was trained on.

Taken together, we can conclude that the geometry of the neuronal representations in RNNs is modulated by the difficulty of the task. Tasks that can be solved by linearly integrating sensory evidence will produce low-dimensional representations (G1; Fig. 7a), whereas more complex tasks that require non-linear integration of evidence will create higher-dimensional representations (G2 and G3). RNNs trained on simple tasks become overly specialized at performing that specific task. As the difficulty of the training task increases, the network could potentially perform a larger number of tasks by simply retraining the output weights. This flexibility comes at the cost of a reduced generalization ability. However, CCGP remains above chance, even for the easy tasks the network has not been trained on. To show that the flexibility is provided by the non-linear components of the responses, we performed an analysis similar to the one of Fig. 7: we fit encoding models with different number of hidden layers to the activity of RNN units (see Methods). Consistently with Fig. 8, we found that the advantage of non-linear encoding models at explaining RNN activity was modulated by the difficulty of the trained task (Supplementary Fig. S18).

These non-linearities greatly enhance flexibility without compromising much the ability to generalize. This is probably why S1 is operating in the regime defined by the geometry G2 of Fig. 7a: it could be a result of training on multiple tasks, or a result of evolution, which selected animals to solve a multitude of complex tasks, generalize to unseen conditions and be robust to noise.

## Discussion

Neural responses in sensory areas are often described in terms of receptive fields. This description is useful and predictive of the responses to novel sensory stimuli only if the responses are linear functions of a relatively small number of stimulus features. This is not the case in somatosensory cortex, in the experiment that we analyzed, in which we observed that non-linear models are the best description of the neural responses. Neurons non-linearly mix multiple variables at different times, leading to an interesting form of spatio-temporal mixed selectivity. The variables are non-linearly mixed even when the task can be solved by simply computing a weighted sum of them (linear readout). Non-linear mixing can easily lead to very diverse neuronal responses as each neuron can respond to a different non-linear function of the stimulus features. Indeed, we observe that the neurons exhibit very diverse responses. However, this does not necessarily mean that the responses are completely disorganized. When one considers the responses of multiple neurons, it is possible to identify an interesting structure in the neural activity space: whisker contacts, which are important features for performing the shape discrimination task, are represented in subspaces that are approximately orthogonal. This kind of factorized or disentangled representations have been observed in the prefrontal cortex [4, 25], in the hippocampus [4], in infero-temporal cortex and in perirhinal cortex [26, 27, 19] and in the motor cortex of monkeys [28], and they are known to have important computational properties for generalization [17, 18].

### Why are the neuronal responses non-linear when the task is linear?

As the non-linearity has a cost in terms of robustness to noise, why are the representations nonlinear even if the non-linearity is not needed in the shape discrimination task? We showed that the observed geometry actually represents a good compromise between the ability to perform complex discrimination tasks, which is typical of high-dimensional representations, and the robustness to noise and the ability to generalize to novel situations, which is typical of low-dimensional representations. This interesting compromise can be reproduced in a simulation of a RNN trained to perform a variety of different tasks that are similar to the shape discrimination task. In these simulations we also observed that the non-linear component of the representations is progressively more important in more complex tasks, and that its cost in terms of noise robustness is relatively small compared to the increase in flexibility. This suggests that somatosensory cortex is operating in an interesting regime that is probably the result of training on a variety of tasks and allows for flexibility and generalization.

### Using encoding models to characterize the collective response properties of populations of neurons

Previous studies showed that the dimensionality of neural representations can be maximal (monkey PFC [5]), very high (rodent visual cortex [16]), or as high as it can be given the structure of the task [29]. More recently, in [4] the authors showed that representations can have the maximal dimensionality required to separate all possible groups of stimuli with a linear decoder (shattering dimensionality) and, at the same time, exhibit a low-dimensional scaffold which allows for cross-condition generalization. All these studies focused on computationally relevant properties of the geometry of the representations of neural populations, ignoring the detailed information about the response of individual neurons. Other studies looked more closely at the components of neuronal responses that are important for characterizing this geometry. Some of these studies focus on the two important ingredients for getting high dimensionality: mixing and diversity [30, 31]. Some others revealed that the responses of individual neurons can be well described by linear mixed selectivity [32, 26, 19], indicating that the representations are low-dimensional, or disentangled, to use the machine learning language [18].

Here we adopted a new approach to characterize both collective properties of the representations and the dynamic response of individual neurons. Indeed, using the neural network encoding models we could characterize the response of an entire population of neurons. This is more than reproducing the responses of all the individual neurons of the population, because the encoding models can capture also the correlations between the activities of different neurons, which can be important to determine the geometry of the representations and their effects on information and behavior (see e.g. [33, 34, 35, 36, 37]). This approach was motivated by the fact that the task involves some form of active sensing, which is closer to natural behavior but far more difficult to analyze. In contrast to the monkey experiments mentioned above [30], the trial temporal structure is highly variable as it depends on the way the animal decides to move the whiskers. Because the sensory input in this kind of behavior involves a larger set of variables which are also continuous, we used a more unbiased approach to identify those variables that could be important to predict the behavior and the neural activity of the animal. We started from this larger set of variables that characterize complex spatio-temporal patterns and let the encoding model find those that are most important.

### Mixed selectivity is defined with respect to a set of variables

One important issue to be discussed is that mixed selectivity is always defined with respect to a set of variables [38]. These variables typically characterize the sensory input and the motor output. Also in our case non-linearities are defined with respect to a set of variables, and this is one of the limitations of all the analyses that focuses on the responses of invidual neurons. For example, neurons could respond to a non-linear combination of two variables *x* and *y*, say *xy*. If one considers *z* = *xy* as an additional variable, then the non-linearity disappears, as the neuron can all be described as a linear combination of *x,y,z*. However, it is important to stress that in our analysis we used the same variables to characterize the task and the neural activity. The non-trivial result is that a linear combination of these variables is the best predictor of the stimulus identity and the choice of the animal, but not of the neural activity. Even when we considered additional variables (e.g. whisker angles were used to predict the activity but not the stimulus or the behavior), we still needed nonlinear interactions. This is significant because these additional variables could be related to non-linear interactions between other variables. Nevertheless, a linear encoding model that has access to all these variables still performs worse than a non-linear one.

### Non-linear spatio-temporal mixed selectivity and the architecture of the neural circuit

Our analysis showed that non-linear interactions are an important component of the neuronal responses. The strongest interactions are between the variables representing whisker contacts and those representing the angle of the whiskers. Interestingly, these interactions are between the angle at the current time and the whisker contacts that occurred in the preceding 100 ms time step. Also the interactions between contacts of different whiskers affect the current neural activity if the contacts happened in the preceding time bin. Although the interactions are delayed, the information about the whisker contacts is not, and the strongest contribution to the neural activity comes from the contacts that happen in the current time step (Figure 5). This means that the first information to arrive in the somatosensory cortex is more linear, and the interaction terms affect the neural activity with a delay of the order of 100 ms. Although it is difficult to draw conclusions, we can speculate that the whisker contact information arrives first from segregated inputs that contain information about separate whiskers. The interaction terms, which appear later, could originate from some non-linear recurrent neural circuit that might be local, within somatosensory cortex, or long-range, involving other areas such as secondary somatosensory, motor, frontal cortex or secondary thalamic nuclei (e.g., [39, 40, 41]). For the information about whisker position (expressed as angles in our analysis) the dominant interaction terms are instead between angles at the current time. It is possible that this information is already non-linearly mixed in other brain areas (downstream, like motor cortex, or upstream like primary thalamus and brainstem).

We also found that all columns in S1 represent information about all whiskers in approximately orthogonal spaces. This result is an important follow-up to [3], in which we did not observe any somatotopic organization. We also do not see significant differences between column selectivity, but our new analysis reveals that there is another interesting organization, as whisker contacts are represented in approximately orthogonal subspaces. This organization is preserved across columns, as much as the non-linear mixing we observed for the responses of individual neurons.

### Non-linearities in artificial neural networks used to model the biological brain

Recently, it has been shown that neural network models (Deep Convolutional Networks, DCN) provide us thus far with the best description of the neuronal responses in the primate’s visual system [23, 24, 42]. All these models, which are constructed by training the networks to perform a classification task, include non-linearities, which certainly play a fundamental role [43, 44]. These results showed also that the classical concept of a receptive field, which is predictive of the responses only when the responses are linear, must be revisited [45, 46]. Interestingly, this approach has even been validated by generating synthetic visual stimuli that are able to maximally activate real neurons [47, 48]. Similarly, accurate models of the somatosensory system could be used to synthesize artificial somatosensory stimuli to drive neuronal activity or even behavior.

### A new method for analyzing experiments that involve real world tasks

Our general framework for analyzing behavioral and electrophysiological data is particularly valuable in experiments in which the animals perform natural tasks, which are becoming increasingly popular in the field [49, 50, 51, 52]. On the one hand, fitting neural networks to predict stimulus identity and animal choice from features extracted from high-speed videos can be extremely useful to identify the most important variables of the task and behavior, respectively. Naturalistic behavior comes with a reduction in the ability to control the strategies followed by the animals, and our approach can potentially provide information about the task and the behavioral strategies for a variety of tasks and animal models. On the other hand, using neural networks to fit neuronal activity from the recorded task variables can be understood as an unbiased multi-dimensional generalization of a population tuning curve. Even though the tuning information is implicitly contained in the architecture and weights of the encoding model, it can still provide crucial insights about the coding properties and geometrical structure of the recorded neuronal population. In our case the animals actively sample the objects by moving the whiskers, and this can greatly complicate the study of the geometry of the neural representations. For example, some of the quantities used in the past to characterize the geometry of neural representations like the shattering dimensionality can require lengthy calculations, involving a number of operations that scales exponentially with the number of experimental conditions. This becomes prohibitive in an experiment like the one we analyzed where we need to consider complex spatio-temporal patterns to characterize the sensory input. Our method can still inform us about the geometry of the representations (it considers the activity of a population of neurons), but without incurring such unfavorable scaling. For all these reasons we believe that the method we propose here can be applied to a number of more natural tasks which are becoming progressively more feasible in the neuroscience community.

## Methods

### Behavioral task and recordings

This experiment has been described in detail in [3]. Here we provide a brief summary of the behavioral setup and data acquisition.

Ten head-fixed mice were trained to perform a shape discrimination task in the dark by making contacts with whiskers C0, C1, C2 and C3 (Fig. 2a). On each trial, either a concave or convex shape (custom designed and 3D-printed) was moved within reach of the mouse’s whiskers with a linear actuator. All trials started at *t* = −2 seconds when the shapes started moving. Shapes were moved with the same speed in all trials and they could stop at three different locations: far, medium and close, which occurred at *t* = −0.9,−0.7 and −0.5 seconds, respectively. Including three different final positions was important to prevent animals from using simpler strategies based on distance to the shape and to force them to integrate contacts across whiskers and time to perform the discrimination task. All trials had a fixed duration of 2 seconds. At *t* = 0 the response window opened and mice had to report their choice by licking either on the left or right lickpipe for concave and convex shapes, respectively. Licks were monitored by infrared beams or capacitive touch sensors. Even though mice were free to lick throughout the trial, the choice on each trial was determined by the side of the first lick after the response window opened (*t* = 0).

Whisker and shape position were recorded with a high-speed camera (200 frames/second). Whisker tracking was based on a modified version of ‘pose-tensorflow’ package [21, 20], which is the ‘feature detector’ network used in the first version of DeepLabCut [22]. The network was trained to track eight equally spaced joints per whisker. Whisker contacts were identified when the distance between the tip of a particular whisker and the edge of the shape was smaller than 10 pixels. Angular position was defined as the angle of the line between the tip and the base of each whisker.

Populations of individual neurons (single units) were simultaneously recorded in mouse somatosensory cortex (S1) during the whisker-based shape discrimination task (Fig. 4a). Mice were implanted with a custom-designed stainless steel headplate between postnatal day 90 and 180. We removed the scalp and fascia covering the dorsal surface of the skull and positioned the headplate over the skull and affixed it. To permit electrophysiological recording we used a dental drill to thin the cement and skull over S1, rendering it optically transparent, and coated it with cyanoacrylate glue. We used intrinsic optical signal imaging to locate the cortical columns of the barrel field corresponding to the whiskers on the face. We then used a scalpel to cut a small craniotomy directly over the columns of interest. Between recording sessions, the craniotomy was sealed with silicone gel. To record neural activity, we head-fixed the mouse in the behavioral arena. We lowered an electrode array using a motorized micromanipulator. We used an OpenEphys acquisition system with two digital headstages to record 64 channels of neural data at 30 kHz at the widest possible bandwidth (1 Hz to 7.5 kHz). We used KiloSort [53] to detect spikes and to assign them to putative single units. We identified inhibitory neurons from their waveform half-width, i.e. the time between maximum negativity and return to baseline on the channel where this waveform had highest power. Neurons with a half-width below 0.3 ms were deemed narrow-spiking and putatively inhibitory. We measured the laminar location of each neuron based on the manipulator depth and the channel on which the waveform had greatest RMS power.

A total of 584 neurons were recorded from 23 sessions that included 7 different mice. The mean number of simultaneously recorded neurons was 25.4. From these 584 neurons, 68 were recorded in layer 2/3, 157 in layer 4, 249 in layer 5 and 96 in layer 6. Also, from the total number of neurons 16% were categorized as inhibitory and 84% as excitatory neurons. All experiments were conducted under the supervision and approval of the Columbia University Institutional Animal Care and Use Committee.

### Decoding of Behavior

On each trial, we built a matrix that contained behaviorally relevant variables through time. In the following, we will refer to this matrix as the spatio-temporal whisking pattern gathered by the behaving mice. We used 20 time bins per feature after dividing 2 seconds into time bins of 100 ms. For whiskers C0, C1, C2 and C3 we included number of contacts and angle of contact (Fig. 3a), since these were shown to be the most informative whisker features for both decoding shape and lick side [3]. In main text and figures, we will use C0, C1, C2 and C3 when referring to whisker identity, and *C*_0_, *C*_1_, *C*_2_ and *C*_3_ when referring to the contacts made by each of these whiskers. The total amount of features on each trial was 160, 8 whisker features (contacts and angle of contacts for each whisker) times 20 time bins. All features for each individual session were normalized to null mean and unit standard deviation. For each mouse, we concatenated all recording sessions into a single super-session, which significantly increased the number of trials used to fit each model. Trials that did not register any lick within the first 500 ms after response time (*t* = 0) were discarded from the analysis. In total we used 10 mice, with a mean of 1266 trials per mouse (super-sessions). All analysis were performed with custom written python and pytorch scripts.

We decoded the identity of the presented shape (stimulus; green) or lick side (choice; blue) on a trial-by-trial basis. In Fig. 3b,c the model was trained after balancing correct and incorrect trials and the quantity to be decoded (stimulus or choice). For instance, when the decoder was trained to predict stimulus identity, we randomly sampled (without replacement) trials from the train set such that correct, incorrect, concave shape and convex shape trials were equally populated. By balancing correct and incorrect trials we ensured that stimulus and choice were uncorrelated. Otherwise, information about choice would have been artificially boosted by stimulus information. We refer to this balancing as decorrelation, and it was repeated 10 times. In Fig. 3b,c the data was split into train, test and validation (2 nested KFold, *k* = 4) in order to optimize the *l*2 regularization strength over the range [10^−7^, 10^3^] (20 steps log-evenly spaced). The reported decoding performances corresponds to the mean across cross-validations and decorrelations on the validation set after optimizing regularization strength on the test set. In Fig. 3b,c we used logistic regression (sklearn).

For Fig. 3b we gradually increased the complexity of the behavioral features to decode stimulus and choice by considering: sum of all contacts across time and whiskers (Sum all), sum of all contacts across whiskers (Sum whisker), sum all contacts across time (Sum time), all contacts across whiskers and time (All contacts) and all contacts and angles of contact across whiskers and time (Contacts + Angle). The inset in 3b corresponds to the weights of the classifier trained after summing contacts across time. Information about stimulus and choice across time was calculated by linearly decoding the cumulative number of features (contacts and angle of contacts) up to that particular time (Fig. 3c).

We analyzed the complexity of the whisker-based shape discrimination task by decoding the spatiotemporal whisking pattern with different decoding models (multilayer feedforward networks with 0, 1, 2 or 3 hidden layers of 100 ReLU units). In the following, because a feedforward network with 0 hidden layers is equivalent to a linear classifier, we will use these two terms synonymously. The models in Fig. 3d were trained and tested following the same steps than for Figs. 3b,c. However, instead of using logistic regression (sklearn) we fit the feedforward networks with stochastic gradient descent (batch size 64, 100 epochs) on pytorch, where the optimal learning rate *η* was obtained following the same procedure than for the regularization strength ([10^−7^,1], 20 steps log-evenly spaced). We used *cross-entropy* loss and ADAM optimizer. The reported decoding performances on correct and error trials (Supplementary Fig. S3a) correspond to the mean performance on the validation set (see above) after splitting trials into correct and error.

For Supplementary Fig. S3b we created two *ad-hoc* tasks from the spatio-temporal whisking patterns gathered by the animals, the easy and the complex tasks. Mice were never trained on these tasks, they correspond to tasks that have been defined on the whisker contact space for whiskers C1, C2 and C3 *a posteriori*. For each mouse we first summed contacts through time on each trial (total contact space). The easy task was defined by splitting the trials in the super-session into two linearly separable classes on the total contact space. For the complex task, trials were split into two non-linearly separable classes (3D-parity) also on the total contact space. In both tasks the contact space was first transformed by a unitary random rotation. Importantly, for both the easy and the complex task, the two classes were equally populated. This was achieved by adding Gaussian noise (standard deviation of 0.1) on each whisker total counts on each trial so that a median split was uniquely defined. Therefore, each trial on a super-session was assigned either to class 1 or class 2 for the easy task, and either to class 1 or class 2 for the complex task. For each mouse, the easy or complex task were performed by reading out the feature matrix that contained whiskers C0, C1, C2 and C3 contacts across time (20 time bins of 0.1 seconds, 80 features in total). The procedure for fitting the different models was the same as in Fig. 3d, with the only difference that we did not need to balance correct and error trials. The *l*2 regularization strength and the learning rate *η* exploration intervals were [10^−6^,1] and [10^−4^,1] respectively, log-evenly spaced in 10 steps. Errorbars in Fig. 3b-d and Supplementary Fig. S3 correspond to the standard error of the mean (s.e.m.) across mice (super-sessions).

### Encoding Models

On each trial we built a matrix that contained all the experimental variables that we considered could affect the firing rate of S1 populations. We analyzed the time interval *t* = [−2.1,1.0] seconds in time bins of 100 ms, which spanned from the beginning of the trial to one second after response window opened (31 time steps per trial). The experimental variables used in the encoding models were: contacts, angle of contact and angular position of whiskers C0, C1, C2 and C3; lick side and lick rate; current and previous reward, stimulus, shape position and choice. We will refer to whisker and lick variables as *continuous-variables* and previous and current reward, stimulus, position and choice as *trial-variables*. For each recording session we concatenated all the time steps across trials (Fig. 4b). On each time step S1 population activity was regressed against the current *continuous-variables* and up to five time steps backwards in time (500 ms = 5 steps ×100 ms). *Trial-variables* were arranged as indicator variables throughout the length of the trial. Population activity was regressed using a total of 70 *continuous-variables* (70 = 14 variables × 5 time steps) plus 248 *trial-variables* (248 = 8 variables × 31 time steps). Both neuronal activity and regressors were normalized to null mean and unit variance. Trials that did not register any lick within the first 500 ms after response time (*t* = 0) were discarded from the analysis. In total we used 23 recording sessions from 7 different mice, with 25.4 mean number of simultaneously recorded neurons and 4883 mean number of effective trials used to fit the models (trials × time steps).

We analyzed the encoding properties of populations of neurons in mouse S1 by regressing the neuronal activity against the experimental variables described above. Similarly to behavior, we used different encoding models with different levels of flexibility (multilayer feedforward networks with 0, 1, 2 or 3 hidden layers of 100 ReLU units). Similar to classification, an encoding model with 0 hidden layers is equivalent to a linear regression. We fit the encoding models by minimizing the mean-squared-error (MSE-loss) between the predicted and the real firing rate (stochastic gradient descent, batch size 64, 100 epochs; Fig. 4). To validate our results with a different loss function, Poisson-loss was also used to fit the models (Supplementary Fig. S6). The linear model can only implement pure and linear mixed selectivity, while encoding models that include at least one hidden layer can implement non-linear mixed selectivity [5, 6]. On each recording session, models were fit by splitting the data into train, test and validation (2 nested KFold, *k* = 4). The partition was performed based on the real trials of the experiment so that time steps from the same trial were always grouped in the same partition. Otherwise, due to the correlation between the neuronal activity on consecutive time steps, performances on the validation set could have been artificially boosted. The optimal regularization strength *l*2 and learning rate *η* were obtained by identifying the values that produced the highest performance on the test set over the ranges [10^−7^,2] and [10^−7^, −1] respectively (20 steps log-evenly spaced). As goodness-of-fit for the different encoding models, we used the metric *R*^2^ = 1 − *Loss/Variance*. The reported *R*^2^ corresponded to the mean across cross-validations on the validation set after optimizing regularization strength and learning rate on the test set. All the encoding models were implemented in pytorch and optimized with the ADAM algorithm. The reported decoding performances on correct and error trials correspond to the mean performance on the validation set (see above) after splitting trials into correct and error (Supplementary Fig. S4b). Errorbars in Fig. 4 correspond to s.e.m. across recorded neurons.

In order to evaluate the individual contributions of regressor (or group of regressors) *x_i_* to the predictability of the population’s firing rate, we evaluated the quantity 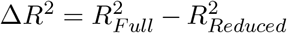 (Fig. 5a), where 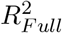 corresponds to the performance of the full model and 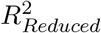 corresponds to the performance of the model when regressor *x_i_* is set to zero. This method is preferred over re-training the whole model without regressor *x_i_* because of the correlations between regressors *x_i_* and *x_j_*, so that we make sure that the reported contribution takes into account the correlation with the rest of regressors.

For each pair of regressors (or pairs of groups of regressors) *x_i_* and *x_j_*, we evaluated the pure nonlinear interaction (contribution) to the encoding model by evaluating 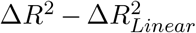 (Fig. 6). Here Δ*R*^2^ corresponds to the loss in predictive power for the non-linear model when both *x_i_* and *x_j_* are set to zero and 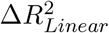 is the equivalent for the linear encoding model. Because non-linear models also include the linear terms, subtracting the contribution from the pure linear model was necessary in order to isolate pure non-linear interactions.

Similar to the synthetic tasks presented in Supplementary Fig. S3b, we also created two synthetic tasks based on whisker contacts: the easy and the complex tasks. Additionally, to benchmark the ability to generalize, we evaluated the cross-condition generalization performance (CCGP). The easy and the complex tasks corresponded to a linear and an XOR task with respect to the contacts of pairs of whiskers, respectively, whereas the CCGP tested how well a linear classifier trained to perform a simple discrimination task on a set of trials would generalize to an unseen set of trials. Given that the encoding models were fit using 5 time steps (100 ms × 5 times steps = 500 ms) for all the *continuous-variables*, for each time step we first summed the number of contacts across the current and previous four time steps for each whisker independently. Gaussian white noise was also introduced in all whisker contacts to obtain a well defined median to create the different tasks (standard deviation of 10^−3^). All tasks were defined as 2D tasks on the summed number of contacts across 5 time steps, so they were constructed from the three different pairs that could be built from the set {C1,C2,C3}: (C1,C2), (C1,C3) and (C2,C3). Importantly, all the time steps in which no whisker contacts were registered for the sum across these 5 time steps were discarded from this analysis. On the easy task, the coloring of the different regions in the whisker contact space (e.g. *C*_1_ vs *C*_3_; Fig. 7b) was defined by a linear boundary, whereas for the complex task it corresponded to an XOR task (Fig. 7c). In both cases the task boundaries were obtained by performing a random unitary rotation on the whisker contact space and splitting each dimension with respect to the median. For the generalization benchmark (CCGP), the process was slightly different. By splitting all the trials into low and high number of contacts for each whisker, we created four different conditions. Cross-condition generalization performance (CCGP; see [4]) was evaluated as the performance of a linear classifier to discriminate between low and high number of contacts for whisker *i* when trained only on low contacts for whisker *j* and tested on high contacts on whisker *j*. For instance, for the (*C*_1_,*C*_3_) pair, a linear classifier was trained to discriminate between low and high number of C1 contacts using only trials of low C3 contacts and tested on high number of C3 contacts (and viceversa). CCGP for whisker C1 corresponded to the mean across training on C2 low and testing on C2 high, training on C2 high and testing on C2 low, and the same process conditioning on whisker C3. CCGPs for whiskers C2 and C3 were evaluated equivalently but conditioning on (C1,C3) and (C1,C2), respectively.

Once the three tasks were defined for each time step, we generated surrogate representations for each encoding model by introducing the pair of whisker contact variables into the different encoding models (Fig. 7e-g). This procedure was only performed on the validation partition. For instance, for (C1,C3) we generated surrogate activity on each time step by introducing in the different encoding models only the experimental variables contacts C1 and C3 for the current and previous four time steps. Given that the final mapping from the last hidden layer of the different encoding models to the surrogate neurons mainly preserves the geometry of the representation (linear transformation), we were able to create new surrogate neurons by generating additional weights from the last hidden layer. In particular, for each set of real weights from the last hidden layer to a particular surrogate neuron, we created extra surrogate neurons by sampling additional weights from a multivariate Gaussian. The mean of this Gaussian was the original set of weights from last layer to that surrogate neuron and the standard deviation was the standard deviation of the real set of weights from the last layer to the original set of surrogate neurons. In other words, from each original surrogate neuron, we created additional neurons with coding properties that were slightly perturbed with respect to the original neuron. From these surrogate representations, linear classifiers (cross-validated logistic-regression) were fit to perform all three tasks. The reported performance in Fig. 7h-j corresponds to the mean performance across cross-validations of the encoding models and pairs of regressors with surrogate populations of 100 neurons. Qualitatively equivalent results were found for surrogate population sizes of {2,10,100,1000,10000} neurons (Supplementary Fig. S14a). The 3D equivalents of the easy and complex tasks and CCGP were also defined from the triplet (*C*_1_,*C*_2_,*C*_3_), and we found qualitatively equivalent results (Supplementary Fig. S14b), even though information for the 3D-complex task (3Dparity) was lower when compared to the 2D-complex task (XOR).

We also evaluated contact information for the different whiskers and columns by decoding whether a set of trials corresponded to a high or low number of contacts for each whisker using neurons recorded from only a particular column of S1 (Supplementary Fig. S13a). CCGPs for the different columns and whiskers was evaluated following the same process described above (Supplementary Fig. S13b). Given that different population sizes were recorded for the different columns of the S1, all the performances in Supplementary Figs. S13 were obtained using 10 neurons, the minimal amount of simultaneously recorded neurons across all columns. We also created the easy and the complex tasks, and the generalization benchmark from other task variables and followed the same procedure described above. In particular, we used the variable continuous angle position of whisker C1, C2 and C3 {*θ*_*C*1_,*θ*_*C*2_,*θ*_*C*3_} (Supplementary Fig. S15a) and the interactions between whisker contacts and continuous angular position (Supplementary Fig. S15b). For Supplementary Fig. S15a the set of pairs (*θ*_*C*1_,*θ*_*C*2_), (*θ*_*C*1_,*θ*_*C*3_) and (*θ*_*C*2_,*θ*_*C*3_) was used, whereas for Supplementary Fig. S15b the interaction terms (*θ*_*C*1_, *C*_1_), (*θ*_*C*1_,*C*_2_), (*θ*_*C*1_,*C*_3_), (*θ*_*C*2_,*C*_1_) (*θ*_*C*2_,*C*_2_) (*θ*_*C*2_,*C*_1_), (*θ*_*C*3_,*C*_1_), (*θ*_*C*3_,*C*_2_) and (*θ*_*C*3_,*C*_3_) were used. Errorbars in Fig. 7, Supplementary Figs. S13, S14 and S15 correspond to s.e.m. across recording sessions.

### Population Decoding

Populations of mouse S1 neurons were recorded during the whisker-based shape discrimination task. Linear classifiers were fit to predict different experimental variables on a trial-by-trial basis. Information about a particular variable for a given time step was calculated using the entire population activity from the beginning of the trial to that particular moment (Figs. 4b). Time bins of 200 ms were used and population activity was normalized to null mean and unit variance. Trials that did not register any lick within the first 500 ms after response time (*t* = 0) were discarded from the analysis. The mean number of simultaneously recorded neurons and trials per session was 25.4 and 157.5, respectively. In total 23 recording sessions from 7 different mice were analyzed. In all panels the data was split into train, test and validation (2 nested KFold, *k* = 4) in order to optimize the *l*2 regularization strength over the range [10^−4^,10^4^] (10 steps log-evenly spaced). In all cases, logistic regression was used as our linear classification model (sklearn).

Shape identity (stimulus; green) and lick side (choice; blue) were predicted on each trial by reading out the population activity (left panel in Fig. 4b). Similar to decoding from the spatio-temporal pattern of whisker features, the classifiers were trained after balancing correct and incorrect trials and the quantity to be decoded (stimulus or choice). We refer to this balancing as decorrelation, and it was repeated 10 times. The reported decoding performances correspond to the mean across crossvalidations and decorrelations on the validation set after optimizing regularization strength on the test set. From population activity we also decoded whether a particular trial corresponded to a high or low number of contacts for the different whiskers (right panel in Fig. 4b). For each whisker we summed the total number of contacts made up to a particular point in time and labeled each trial according to whether it was below or above the median number of contacts. Gaussian noise was added in all trials (standard deviation of 0.1) to obtain a unique median. The reported decoding performances correspond to the mean across cross-validations on the validation set after optimizing regularization strength on the test set.

Populations of recorded neurons were also used to perform the easy and the complex tasks and the generalization benchmark (brown bars; Fig. 7h-j) (see previous section). From all the neuronal recordings, pseudopopulations of neurons were constructed and linear classifiers (logistic regression) were fit to perform these three tasks. To define the easy and complex tasks and the generalization benchmark (CCGP) on each recording session, we first summed the number of contacts throughout the entire trial (2 sec). Similar to the equivalent analysis on surrogate representations (see previous section), all three analysis were defined with respect to pairs of whisker contacts variables: (*C*_1_,*C*_2_), (*C*_1_,*C*_3_) and (*C*_2_,*C*_3_). For the easy and complex tasks, a random unitary rotation was performed on the whisker contact space for a given pair and all those trials that did not include whisker contacts were discarded from the analysis. Four experimental conditions corresponding to low and high number of contacts for two whisker variables were defined. From each experimental condition, 400 and 100 trials were sub-sampled with replacement for the train and test set, respectively. The simultaneously recorded activity of S1 neurons across a particular trial was flattened with respect to the time axis (200ms time bins; 10 time bins per trial). For a given recording session we constructed a train matrix with dimensions 1600 (400 trials per condition × 4 experimental conditions) and number of neurons × 10 time bins, and a test matrix with dimensions 400 (100 trials per condition × 4 experimental conditions) and number of neurons × 10 time bins. It is important to note that by this procedure the train and test matrix did not share any trials, which would artificially boost the estimated performance for the different tasks. For each recording session we repeated this procedure and stacked the different train and test matrices along the dimension of neurons. A total of 584 neurons were recorded across all sessions, which produced a train and a test matrix with 5840 columns (584 × 10 time bins). Two different linear classifiers were fit on the train matrix and tested on the test matrix to perform the easy and the complex task, respectively. The reported performances on the easy and complex tasks in Fig. 7h,i (brown bars) corresponds to the mean across 100 iterations of this process.

In order to evaluate the generalization properties of the recorded neurons (CCGP), we proceeded in a similar way but we worked on the original whisker contact space instead (no unitary rotation). Also, given that the cross-validation is performed across conditions when evaluating the CCGP, only one matrix of pseudopopulation activity was constructed by sub-sampling with replacement from each experimental condition (500 trials per condition). Similarly, the reported CCGP in Fig 7j (brown bars) corresponds to the mean across 100 iterations of this process. To evaluate whether the reported CCGPs for the real recordings were significantly above chance, for each panel we constructed a null-hypothesis distribution and evaluated the probability of obtaining the real CCGP when sampling from it. We followed the same procedure described in [4]. In short, each experimental condition was randomly rotated in the activity space by shuffling each trial with respect to the identity of the neurons. The same random shuffle was used for all trials in a given condition. This procedure destroys the geometrical structure of the representation but maintains the distance between the different conditions approximately constant. We performed 1000 iterations and computed the probability of obtaining the real CCGPs. In all cases, P < 10^−3^ corresponded to CCGPs > 0.58 (distribution not shown), which indicates that the reported CCGPs for the real recordings are all well above chance.

### Recurrent Neural Networks

Recurrent neural networks (RNNs) were trained to perform a task similar to the whisker-based shape discrimination task. The recurrent network consisted of 60 ReLU units whose activity at time *t* (**h**_*t*_) was determined by the following equation:

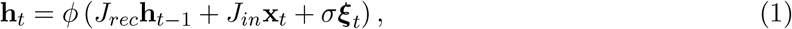

where *ϕ*() is the ReLU non-linearity, ***ξ***_*t*_ is independent and unitary Gaussian noise on each time step and *σ* is the strength of this noise (*σ* = 1 in all our units).

The stimulus **x**_*t*_ consisted of three channels that on each time step could be either 0 or 1, an artificial analogy of whiskers C1, C2 and C3 making contacts or not. On each trial, each input channel corresponded to a random realization of a *Bernoulli* process (*T* time steps) with two possible underlying mean values λ_*low*_ or λ_*high*_. This made a total of 8 different experimental conditions (2 conditions per channel and 3 channels) (Fig. 8). From these 8 experimental conditions we defined three different tasks, the easy, the medium and the complex task. For all tasks, the input information was transformed by an unitary random rotation (same rotation in all time steps and trials). We fit a different RNN for each task, and in each RNN input, recurrent and output weights were trained. The easy task was defined as a task that linearly separated the 8 experimental conditions into 2 groups of 4 (top panel in Fig. 8b); the medium task corresponded to a 2D-XOR with respect to C1 and C2 (top panel in Fig. 8c); and the complex task was defined as a 3D-parity with respect to all channels C1, C2 and C3 simultaneously (top panel in Fig. 8d). Given 8 experimental conditions, there were 3 different easy tasks: separation with respect to the C1 axis only (easy task 1); C2 axis only (easy task 2); and C3 axis only (easy task 3). There were also 3 different medium tasks: separation with respect to a 2D-XOR on (C1,C2) (medium task 1); on (C1,C3) (medium task 2); and on (C2,C3) (medium task 3). There was only one complex task, a 3D-Parity task with respect to all channels. In Fig. 8, easy task, easy other, medium task and complex task corresponded to easy task 1, the mean across easy tasks 2 and 3, medium task 1 and complex tasks, respectively. In Supplementary Fig. S16 all tasks were used.

To recreate the experimental conditions, inputs lasted for 20 time steps but a random delay of Δ*t* = [0, 9] time steps was introduced at the beginning of each trial. All networks were trained to make a decision at *T* = 20. The three networks were trained on datasets of 400 trials per experimental condition and for all channels λ_*low*_ = 0 and λ_*high*_ = 1. We used *cross-entropy* as loss function, the *l*2 regularization strength was set to 10^−10^ and the learning rate *η* = 0.005. We used the ADAM optimizer, batches of 20 trials and as many epochs as necessary to reach 10^−3^ error on the loss function (~ 10 epochs for the easy task, ~ 20 for the medium, and ~ 50 epochs for the complex task). Once trained, networks were tested on 40 trials per experimental condition and for all channels λ_*low*_ = 0.35 and λ_*high*_ = 0.65 for the easy task, λ_*low*_ = 0.3 and λ_*high*_ = 0.7 for the medium task and λ_*low*_ = 0.23 and λ_*high*_ = 0.77 for the complex task.

For each network, input, recurrent and output weights were learned. Additionally, for each network, recurrent and input weights were frozen and readout weights for the other tasks were also trained on the activity of the artificial units (logistic regression). For instance, for the network trained on the easy task (easy task 1) in Fig. 8b (central panel), all learnable weights were optimized for the easy task using backpropagation through time (dark brown). However, additional readout weights on the artificial units’ activity were also trained for the orthogonal other easy task (easy tasks 2 and 3; light brown), the medium task (medium task 1; black curve) and the complex task (maroon curve). These additional readout weights were trained on the train set at decision time (*T* = 20) and tested on the test set on all time steps. In Supplementary Fig. S16 we show the performance curves for additional readout weights when trained on all tasks (easy and medium tasks 1,2,3 and complex task). For Figs. 8c,d (central panel) all weights were trained to perform medium task 1 and complex task, respectively. Similarly, the rest of tasks in Figs. 8c,d were also performed by training a linear classifier (logistic regression) on the artificial units’ activity.

We also evaluated the ability of each network to generalize to unseen experimental conditions by means of the cross-condition generalization performance (CCGP). A very similar procedure to Fig. 7 was used to evaluate CCGP for the three different RNNs. For instance, in Fig. 8b, a linear classifier was trained to perform the easy task (easy task 1; dashed dark brown) by reading out the activity of the artificial units. The classifier was trained on the set of trials defined by easy task 2 = +1 and tested on the set of trials that defined easy task 2 = 0 (and vice-versa). The same procedure was followed for the set of trials defined by easy task 3 = +1 and tested on trials defined by easy task 3 = 0 (and vice-versa). The reported CCGP was the mean across these four procedures. For the rest of CCGPs, the same train-test procedure was followed as defined by the rest of orthogonal easy tasks (see Supplementary Fig. S17).

For each panel in Fig. 8, and Supplementary Figs. S16 and S17 we trained and tested 50 different networks and reported the mean performance across test sets. Each network was trained on a different random realization of the input and rotation. In Supplementary Figs. S16 and S17 the low, medium and high noise levels corresponded to (λ_*low*_ = 0.23, λ_*high*_ = 0.77), (λ_*low*_ = 0.3, λ_*high*_ = 0.7) and (λ_*low*_ = 0.35, λ_*high*_ = 0.65), respectively.

We analyzed the complexity of the different tasks in the same way we analyzed the complexity of the whisker-based shape discrimination task (see Fig. 3). For all trained networks, we used different classifiers with different levels of flexibility (multi-layer feedforward networks with 0, 1, 2 or 3 hidden layers of 100 ReLU units). These classifiers were trained to predict the output of the easy, the medium and the complex task on a trial-by-trial basis by reading out the spatio-temporal pattern that was used as input to the networks (Fig. S18a). The *l*2 regularization strength and the learning parameter *η* were optimized over the ranges [10^−8^,2] and [10^−6^, 0] respectively (10 steps log-evenly spaced). The errorbars for each panel in Fig. S18a correspond to the s.e.m. across 4 different network instances.

The encoding properties of the artificial units on each network were also analyzed in the same way we analyzed the population activity of mouse S1 neurons (Fig. S18b), by fitting encoding models of feedforward networks of 0, 1, 2 and 3 hidden layers. In this case, the *l*2 regularization strength and the learning parameter *η* were optimized over the ranges [10^−8^,10^2^] and [10^−6^,0] respectively (10 steps log-evenly spaced). The errorbars for each panel in Fig. S18b correspond to the s.e.m. across 240 neurons (4 networks × 60 units).

## Acknowledgements

We would like to thank the members of the Center for Theoretical Neuroscience, Marcus K. Benna and Mattia Rigotti for all their insightful comments and suggestions. Support was provided by NINDS/NIH (R01NS094659, R01NS069679, F32NS096819, and U01NS099726); NSF 1707398 (Neuronex); the Gatsby Charitable Foundation (GAT3708); the Simons Foundation; the Swartz Foundation; Northrop Grumman; a Kavli Institute for Brain Science postdoctoral fellowship (to CR); and a Brain and Behavior Research Foundation Young Investigator Award (to CR).

## Author Contributions

RN and SF conceived the project and the analytic approach. CR developed the behavior, videography and performed the electrophysiological recordings. RN analyzed the data. RN, CR, RB, and SF decided how to interpret the results. RN and SF wrote, and CR and RB edited the manuscript.

## Supplementary Information

**Figure S1:**
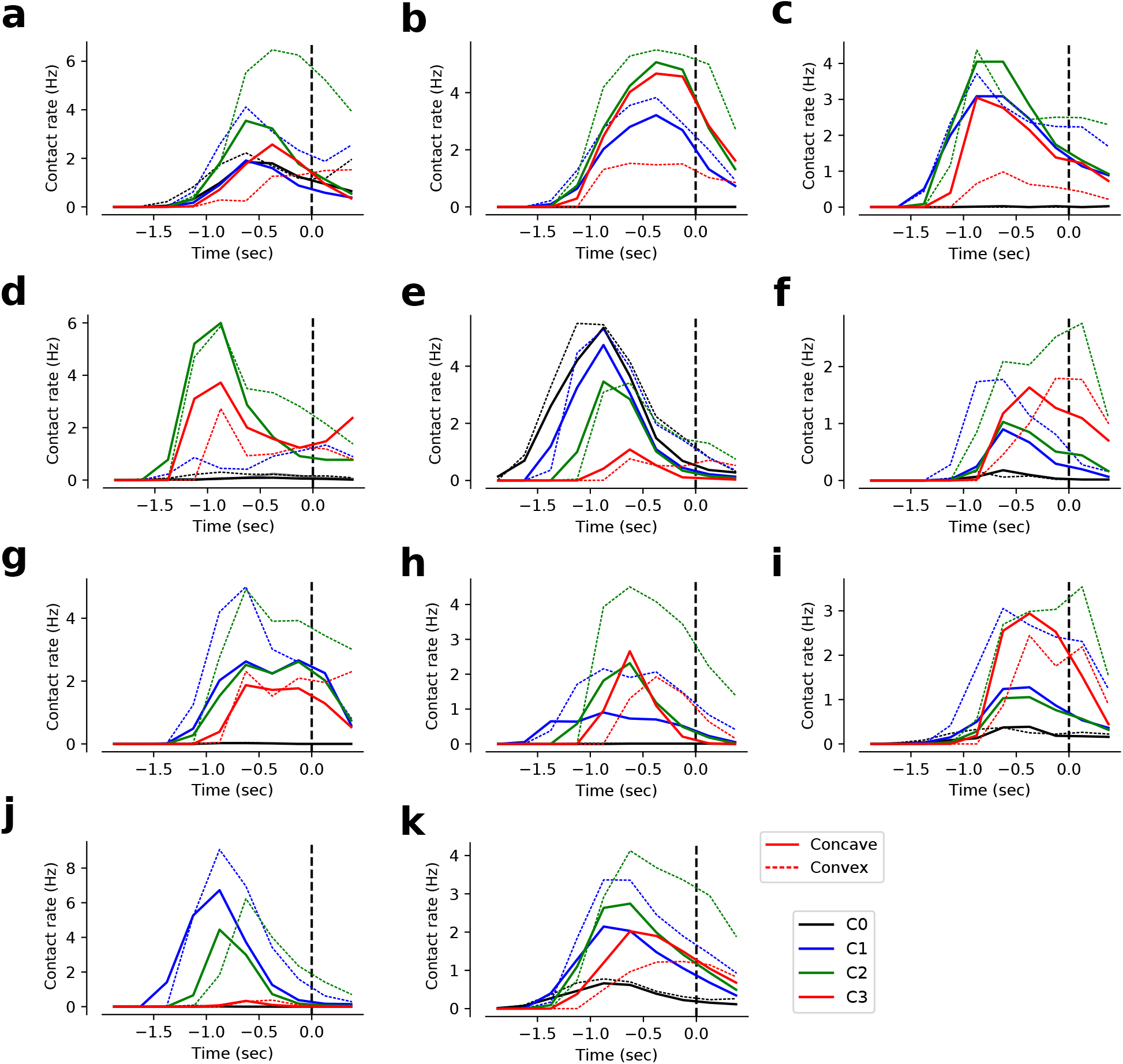
Contact rate (y-axis) as a function of time throughout the trial (x-axis), separately for convex and concave shapes. Contacts were higher for convex than concave shapes for whisker C1 and C2, whereas whisker C3 showed the opposite trend. (**a-j**) Results for all the mice. (**k**) Mean results across mice.

**Figure S2:**
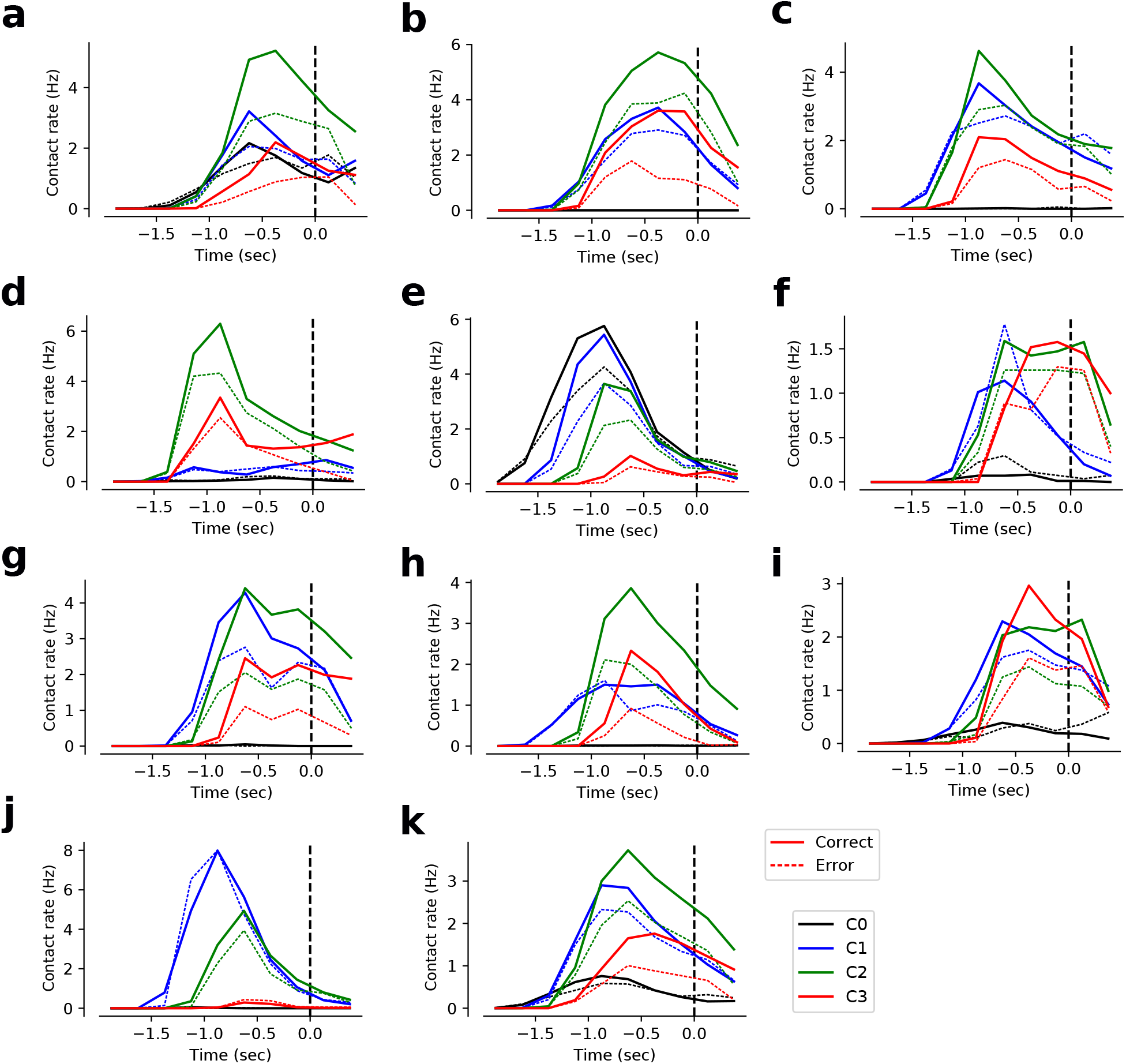
Contact rate (y-axis) as a function of time throughout the trial (x-axis), separately for correct and error trials. Contacts were higher for correct than error trials for all whiskers and animals, (**a-j**) Results for all the mice. (**k**) mean results across mice.

**Figure S3:**
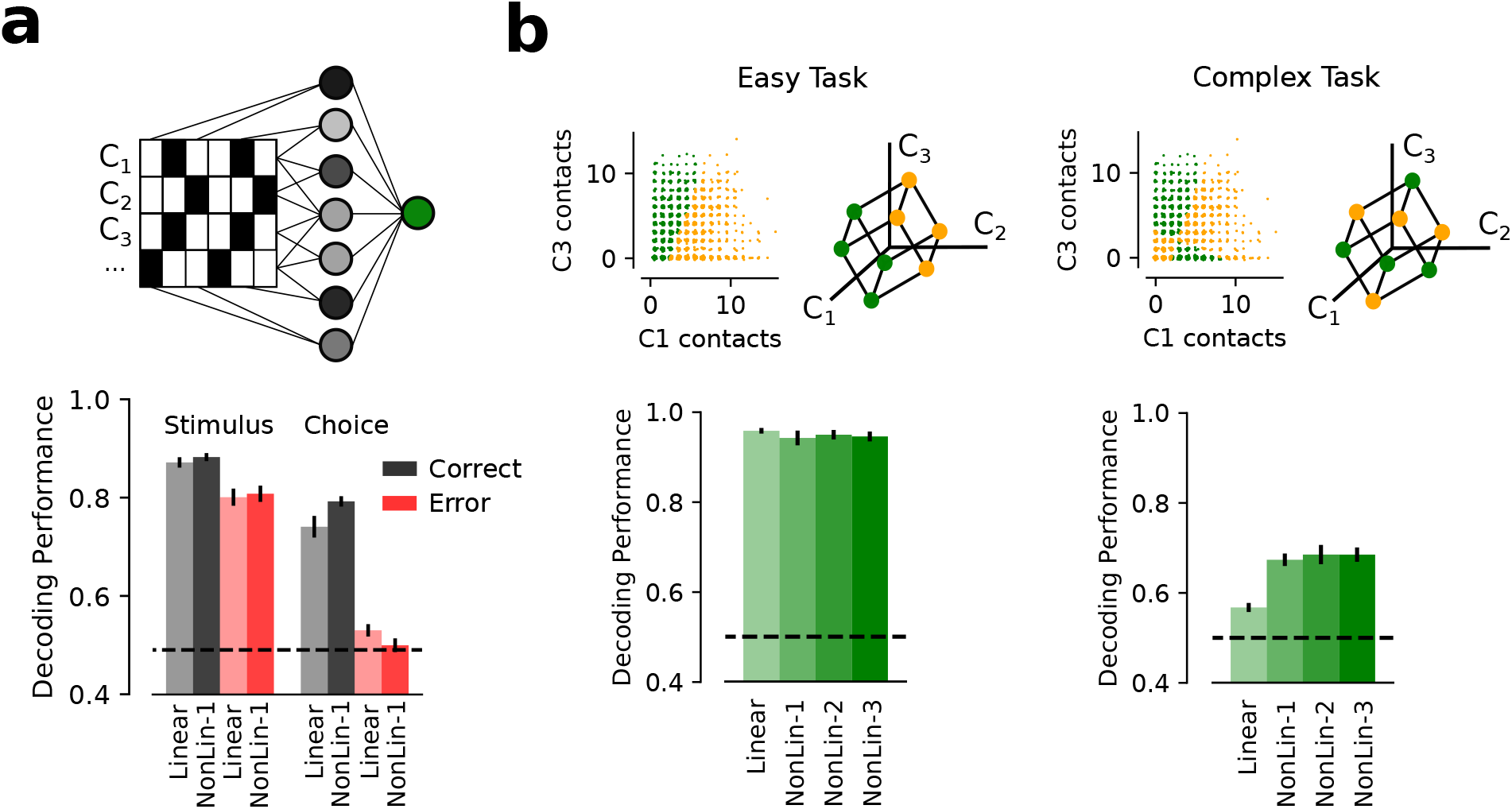
(**a**) A multi-layer feedforward network model is trained to use the full spatio-temporal pattern of contacts and angle of contacts to predict the stimulus and the choice of the animal on a trial-by-trial basis (see Fig. 3). Models were trained using all trials and tested on correct (black) and incorrect (red) trials. Only linear and non-linear models with one hidden layer are shown. Stimulus decoding (left) produced higher decoding performance for correct trials than errors, probably due to the higher number of contacts made by mice on correct trials. On the contrary, correct trials conveyed much more information about animals’ choice than incorrect trials (right). One possible explanation of these effects is that in approximately 60% of the trials animals make very accurate choices that are based on properly sampled sensory cues. In the other 40% of the trials, animals still sample information properly but their choice is inaccurate and based on a hidden variable we do not have access to [54, 55]. (**b**) Decoding performance (y-axis) for the different decoders (x-axis) for the easy (linearly separable; left panel) and complex (non-linearly separable, 3D-parity; right panel) tasks (see Methods). Non-linear cue integration is only advantageous when the task itself requires complex sensory integration across time and whiskers. Error bars in all panels correspond to s.e.m. across mice.

**Figure S4:**
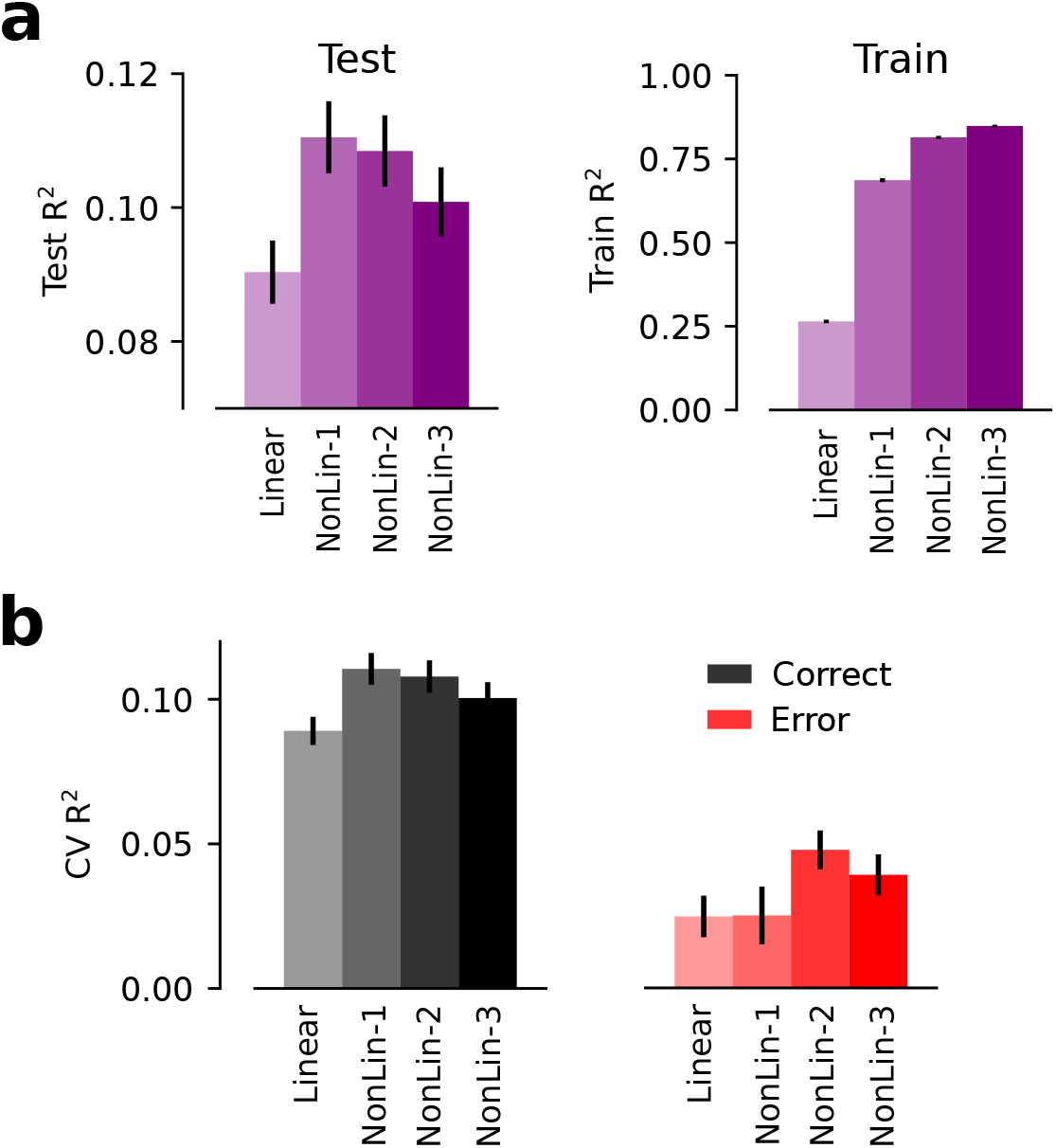
(**a**) Goodness-of-fit (y-axis; *R*^2^) for the different encoding models (x-axis) on held-out data (Test; left) and on the data used for training (Train; right). For held-out data, the best model is a feedforward fully connected network with only one hidden layer (NonLin-1). For the train data, the more complex the model (more parameters), the better the prediction. (**b**) Cross-validated *R*^2^ (y-axis) on populations of S1 neurons for the different encoding models (x-axis) when evaluated in correct (left) and incorrect trials (right). Encoding models explained S1 activity better on correct than error trials (right panel). Errorbars correspond to s.e.m. across neurons.

**Figure S5:**
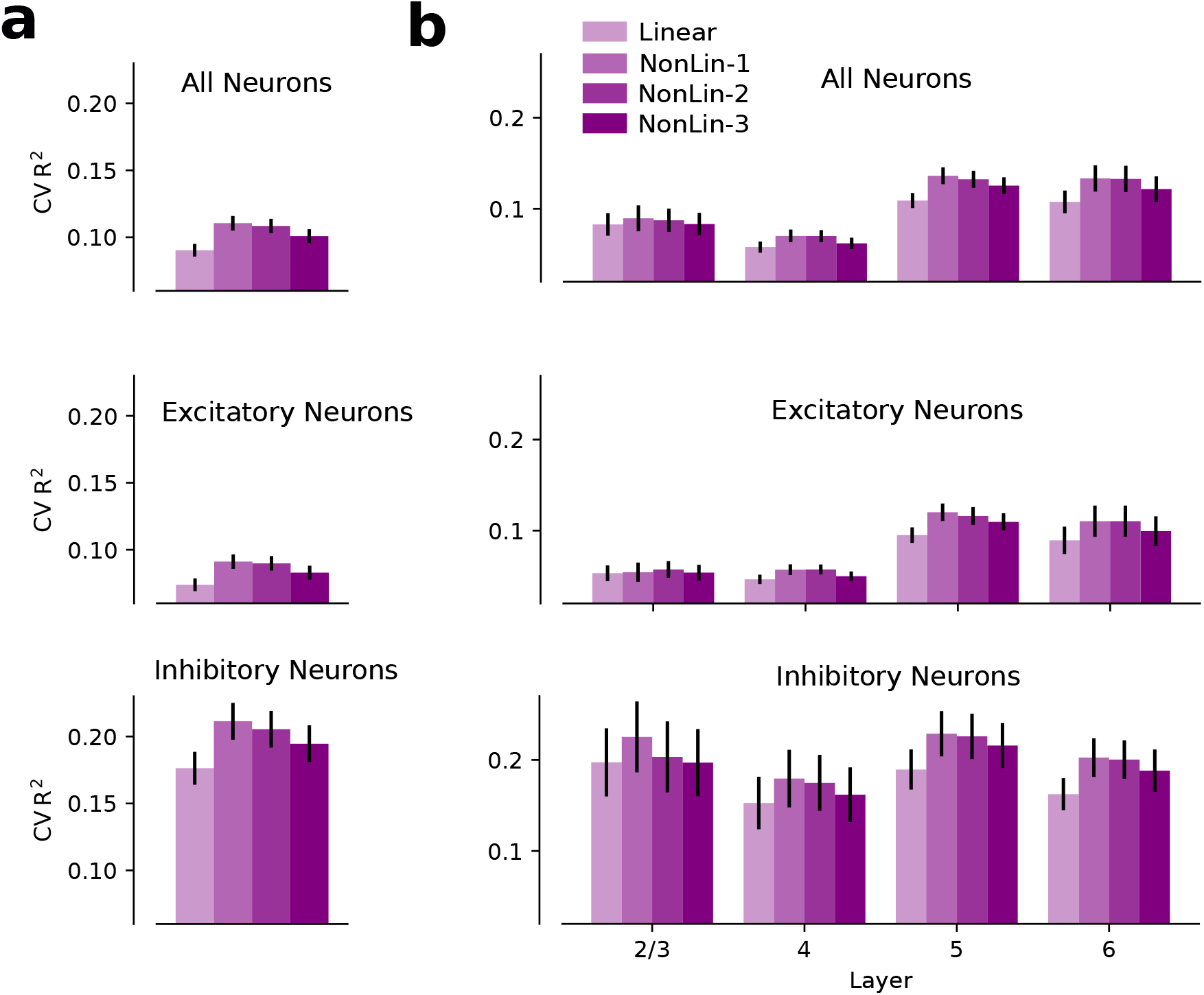
The encoding models explain better the responses in inhibitory neurons and deeper layers. (**a**) Performance (CV *R*^2^) of the different encoding models (x-axis) on held-out data for all neurons (top), only excitatory (middle) and only inhibitory neurons (bottom). (**b**) Performance of the different encoding models on held-out data for neurons across layers for all (top), excitatory (middle) and inhibitory neurons (bottom).

**Figure S6:**
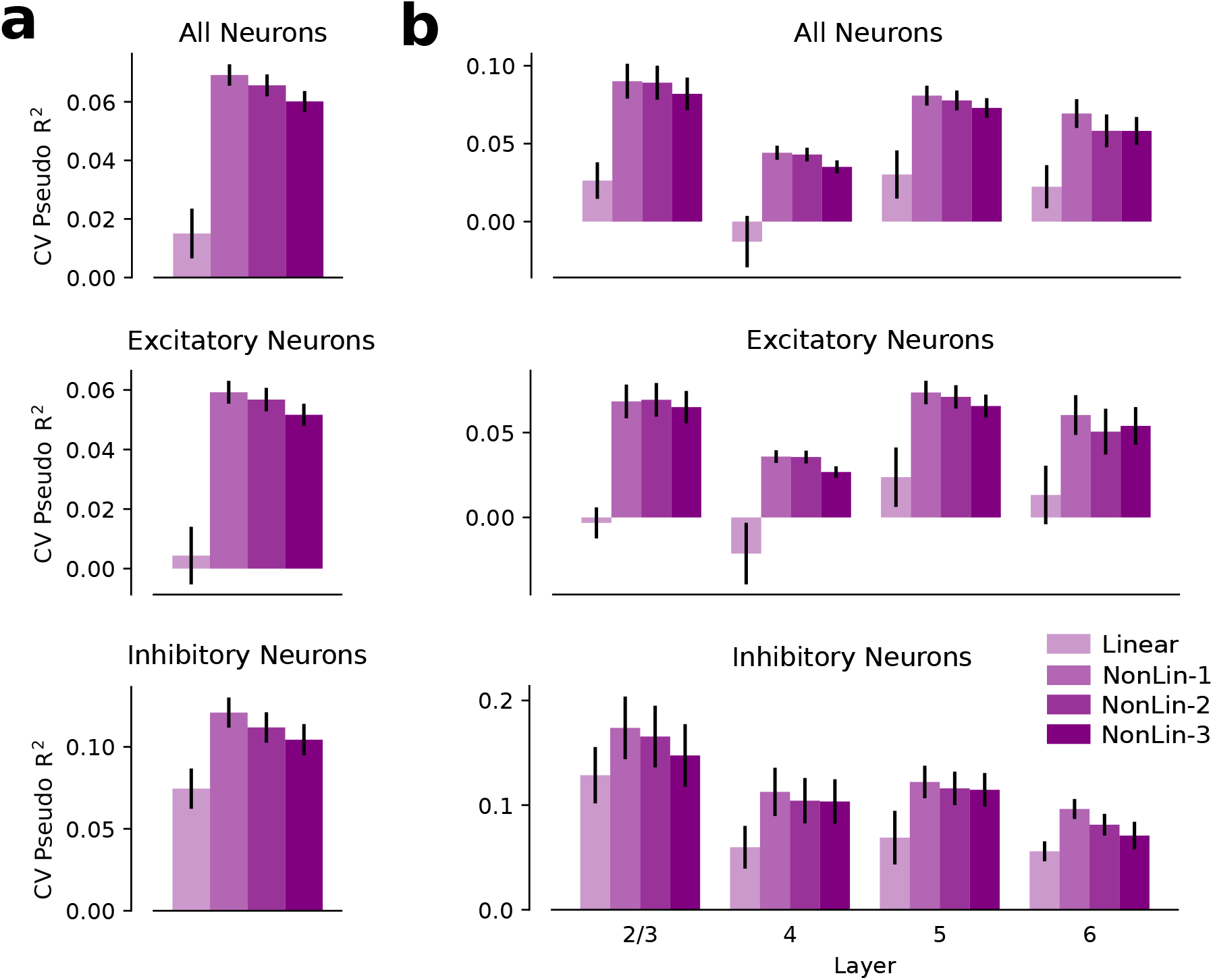
Encoding fits when using Poisson loss instead of mean squared error (MSE). The y-axis shows the Poisson-loss equivalent of the *R*^2^, the Pseudo-*R*^2^. The Pseudo-*R*^2^ is calculated as 1 - PLoss/Variance, where PLoss is the negative Log-likelihood of the Poisson model. (**a-b**) See Supplementary Figure S5.

**Figure S7:**
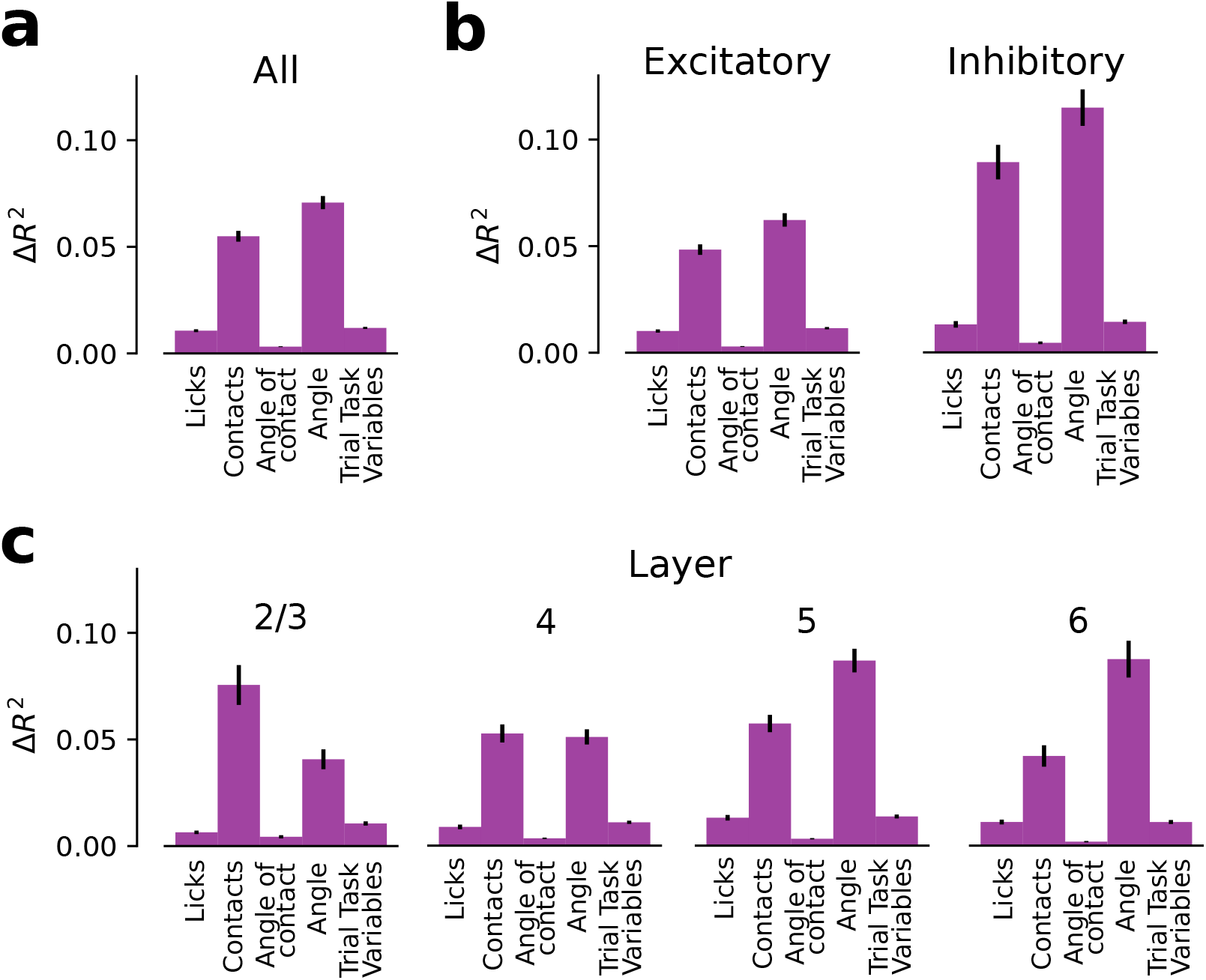
Difference between *R*^2^ of the full model 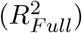 and the *R*^2^ for the different ablations 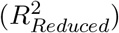 (y-axis; 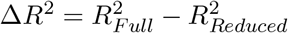). The *R*^2^ for the different ablations (x-axis) is calculated by testing the model on trials where those features have been set to zero. (**a**) Δ*R*^2^ for all neurons. Whisker contacts (Contacts) and continuous angle (Angle) are the most important features in predicting S1 activity. (**b**) Δ*R*^2^ excitatory and inhibitory neurons. Inhibitory populations show a higher Δ*R*^2^ because their *R*^2^ is overall higher (see Supplementary Fig. S5). (**c**) Δ*R*^2^ across layers of the somatosensory cortex. Whisker contacts have a stronger effect on superficial layers (2/3), while whisker angle has a stronger effect on deeper layers.

**Figure S8:**
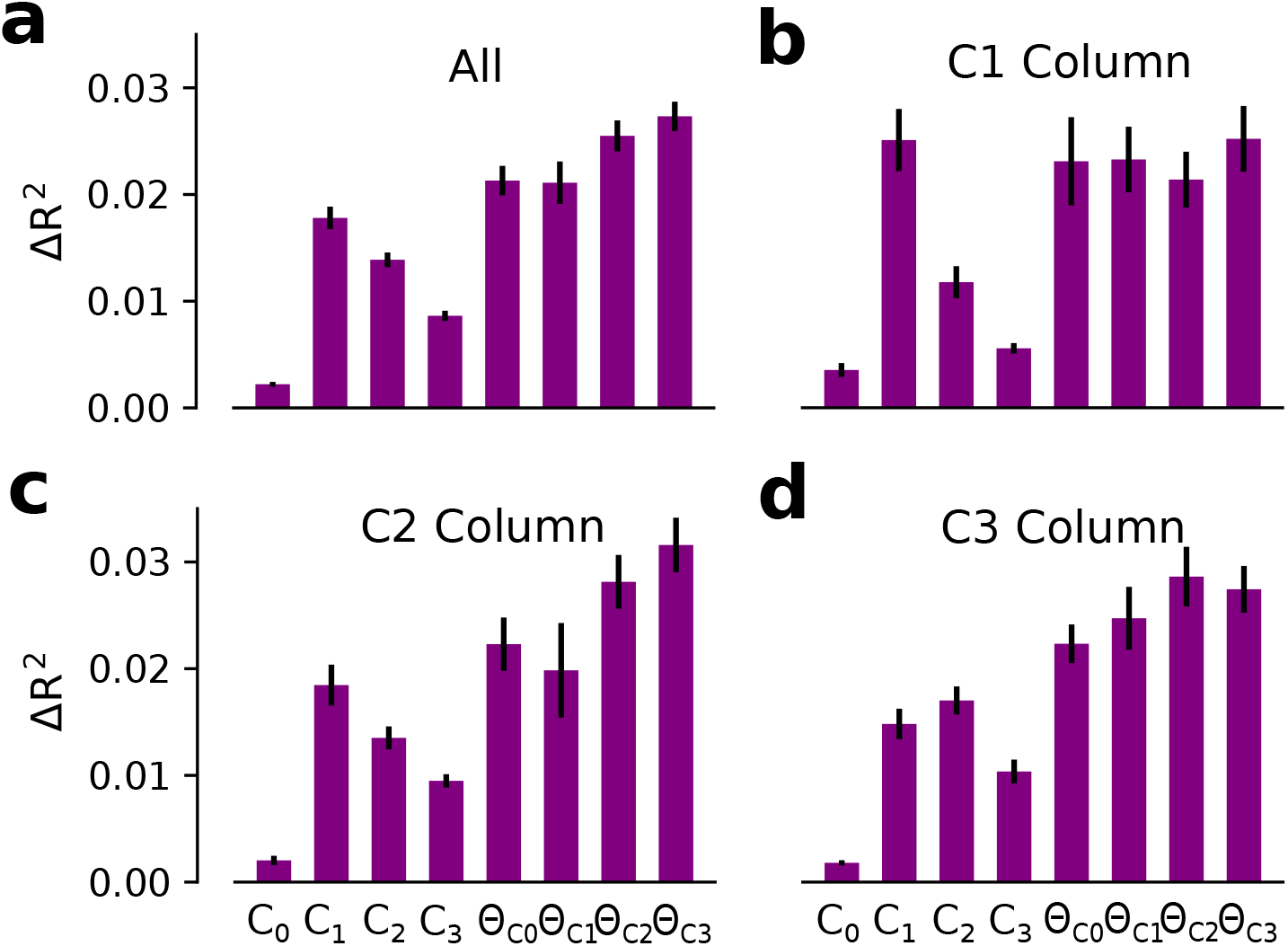
The metric Δ*R*^2^ reveals that neurons in S1 do not strictly obey somatotopy during the whisker-based discrimination task (see also [3]). (**a**) The contribution to the encoding model’s performance (y-axis; Δ*R*^2^) for whiskers’ contacts (*C*_0_, *C*_1_, *C*_2_ and *C*_3_) and angular position (*θ*_*C*0_,*θ*_*C*1_,*θ*_*C*2_, and *θ*_*C*3_) for all neurons. (**b-c**) whisker and angular position encoding strength for C1 column (b), C2 column (c) and C3 column (d). While C1 contacts is the strongest driver in C1 column, C2 and C3 columns are not dominated by C2 and C3 contacts, respectively. Errorbars in all panels correspond to s.e.m. across neurons.

**Figure S9:**
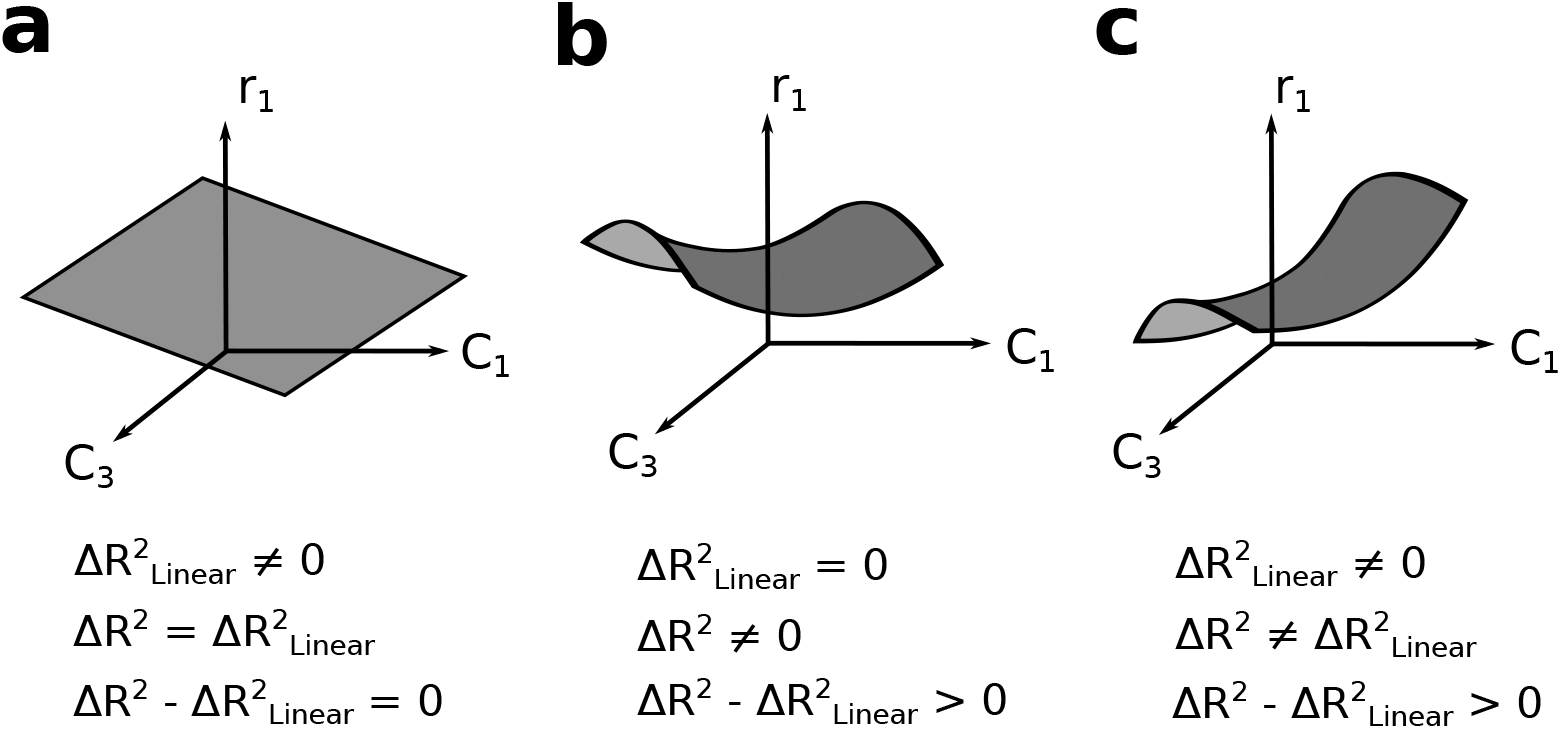
Different coding scenarios would produce different values for 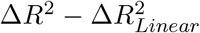. Here we show different tuning schemes with respect to *C*_1_ and *C*_3_ for a fictional neuron (*r*_1_). (**a**) The metric 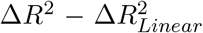 was used to evaluate to what extend the pure non-linear terms were important to predict the population’s firing rate. If the relationship between neuronal activity and encoding variables is linear, 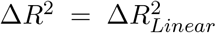 and therefore 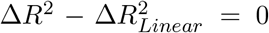 (**b**) If the relationship between neuronal activity and encoding variables is purely non-linear, 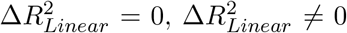 and 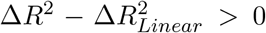. (**c**) If the encoding model is composed of both linear and non-linear components 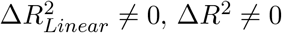, Δ*R*^2^ ≠ 0 and 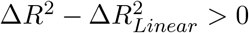.

**Figure S10:**
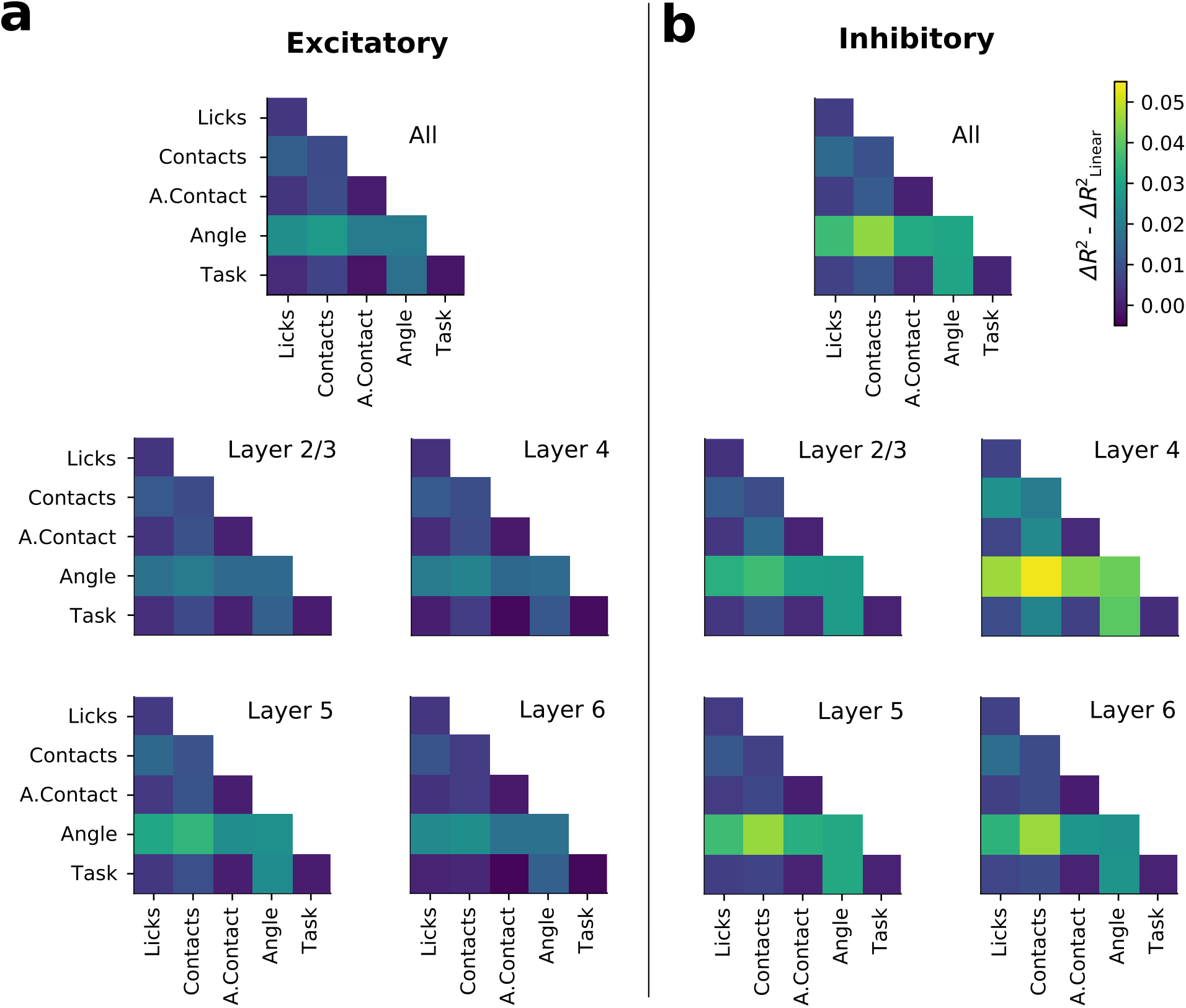
Pure non-linear mixed selectivity contribution 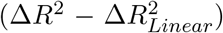 for the interaction between the different blocks of variables across neuronal types and S1 layers. Results were qualitatively equivalent for the excitatory (a) and the inhibitory (b) populations.

**Figure S11:**
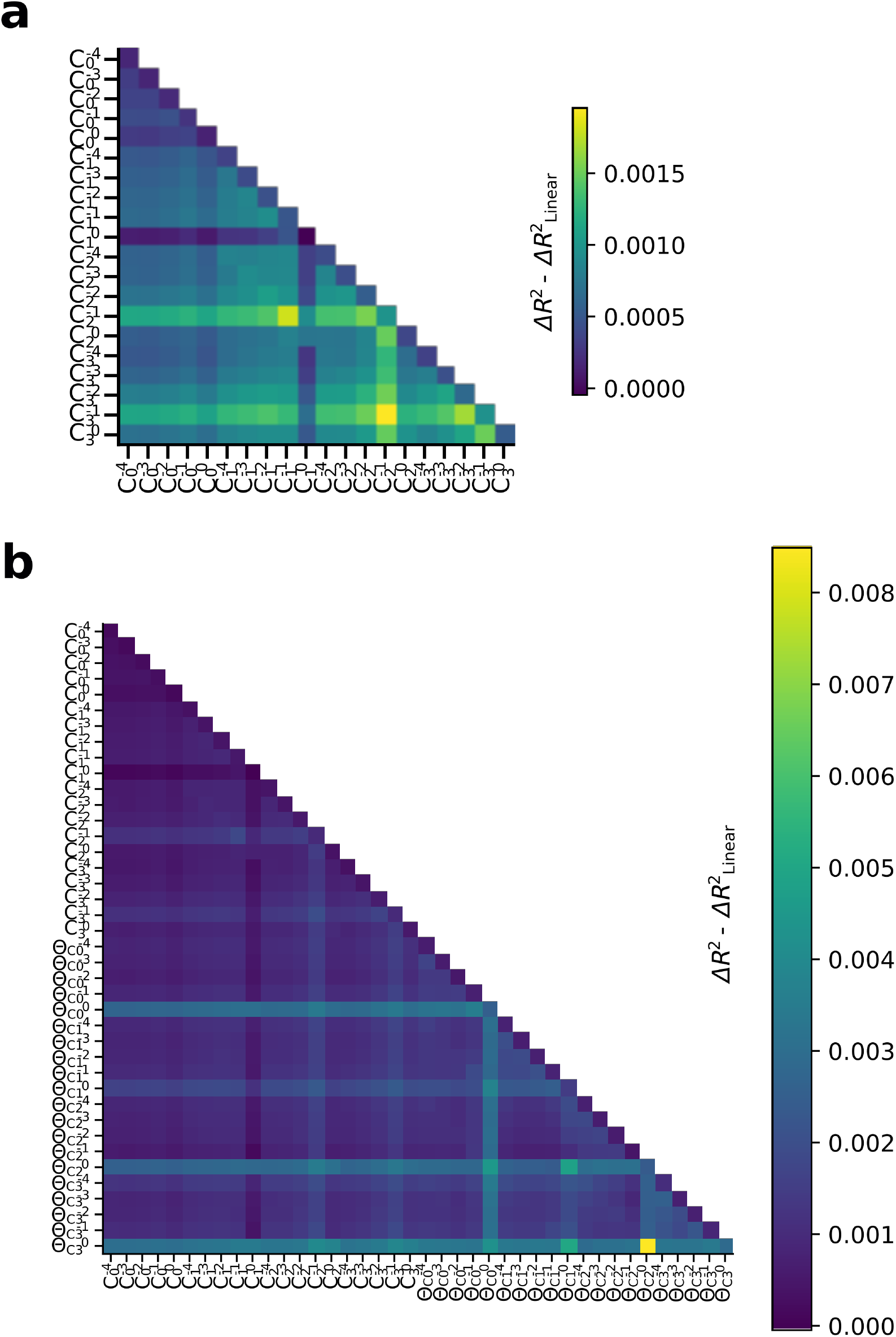
(**a**) Pure non-linear mixed selectivity contribution for the interaction between contacts for the different time steps (time lags) and whiskers 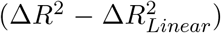. The strongest non-linear contribution in whisker contacts occurs on time lags between neuronal activity and contacts of 100 ms (1 time step) for all whiskers. (**b**) Pure non-linear mixed selectivity contribution 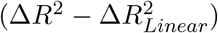 for the interaction between contacts and angular position for the different time steps (time lags) and whiskers. The strongest non-linear contribution in whisker angular position occurs on the current time step for all whiskers. The strongest non-linear contribution for the interaction between contacts and angular position occurs also on time lags between angular position (and neuronal activity) and contacts of 100 ms (1 time step).

**Figure S12:**
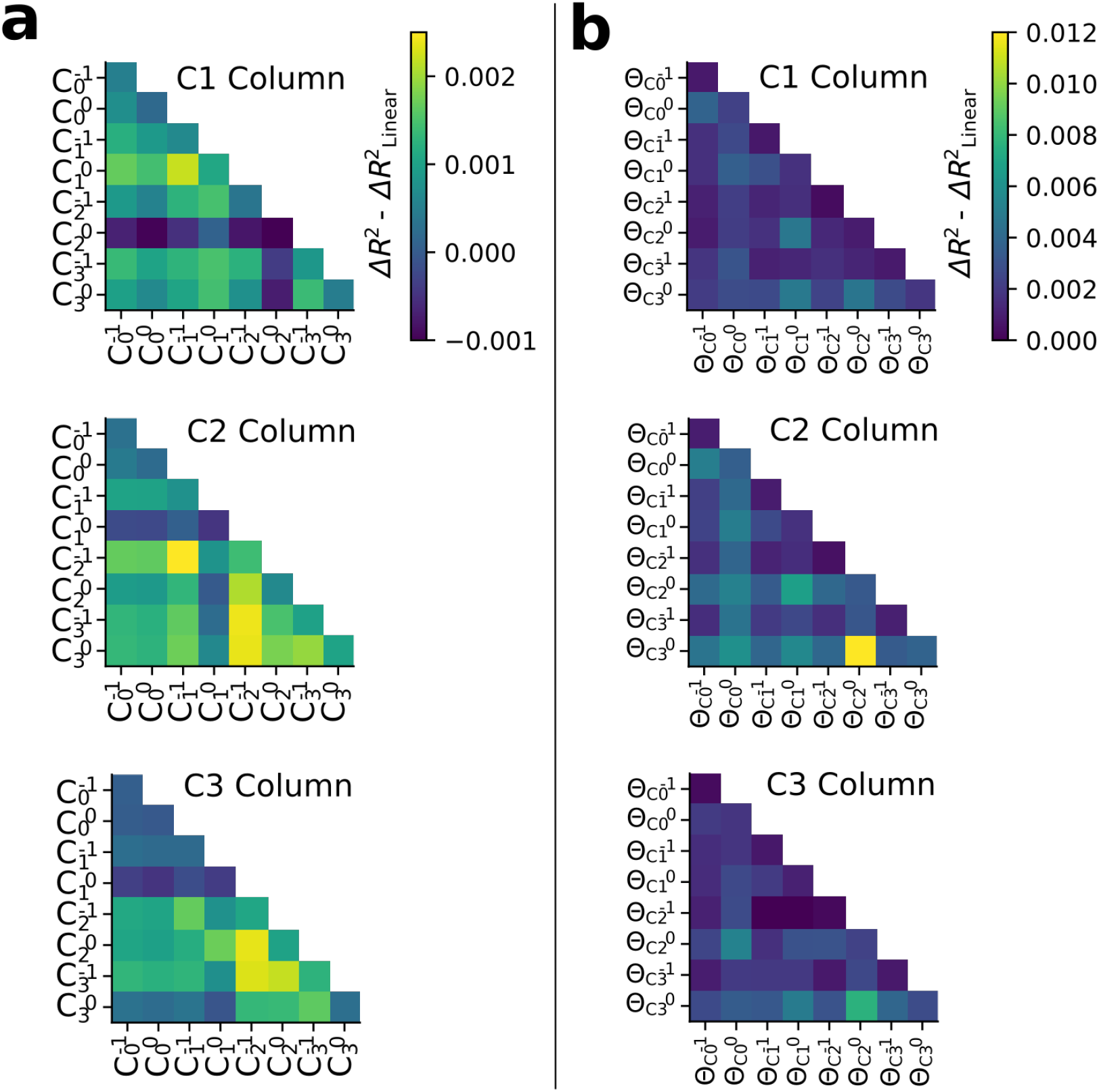
(**a**) Pure non-linear mixed selectivity 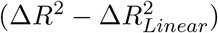 contribution for the interaction between contacts for the different time steps (time lags) and whiskers separately by columnar location of each neuron. The strongest interactions occur at the time lags of 100 ms (1 time step). Even though C1 column shows that C1 terms have the strongest interaction, C2 and C3 columns present a more heterogeneous interaction pattern. (**b**) Pure non-linear mixed selectivity 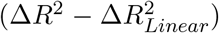 contribution for the interaction between angular position for the different time steps (time lags) and whiskers. The strongest interactions occurs at time lags of 0ms. All columns present strong interactions terms with the rest of whiskers.

**Figure S13:**
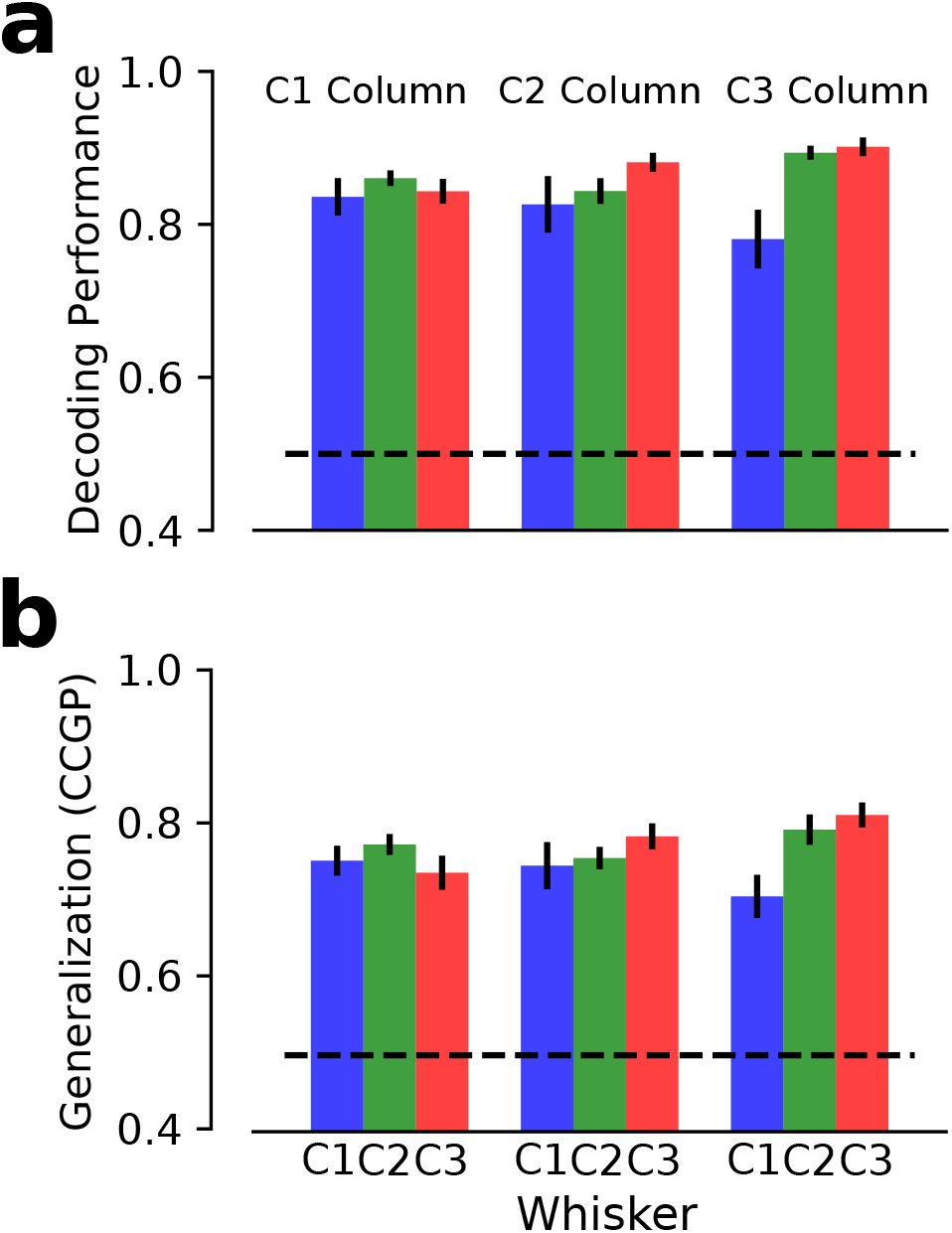
(**a**) Decoding performance on whether the sum of whisker contacts across the current and previous four time steps corresponded to a high or a low number of contacts with respect to the median for each different whisker (C1 blue; C2 green; C3 red) and columns (see Methods). Information about whisker contacts for all whiskers was present in all columns. Decoding Performance was evaluated on surrogate activity generated by the best encoding model (NonLin-1). In each recording session, activity from only one column was recorded. In order to compare information across columns, surrogate activity was generated for 10 neurons, which corresponded to the smallest number of simultaneously recorded neurons across recording sessions (columns). (**b**) Qualitatively equivalent results were found when the CCGP was evaluated (see Methods). All columns encode information about all whiskers in approximately orthogonal spaces. Errorbars in all panels correspond to s.e.m. across recording sessions.

**Figure S14:**
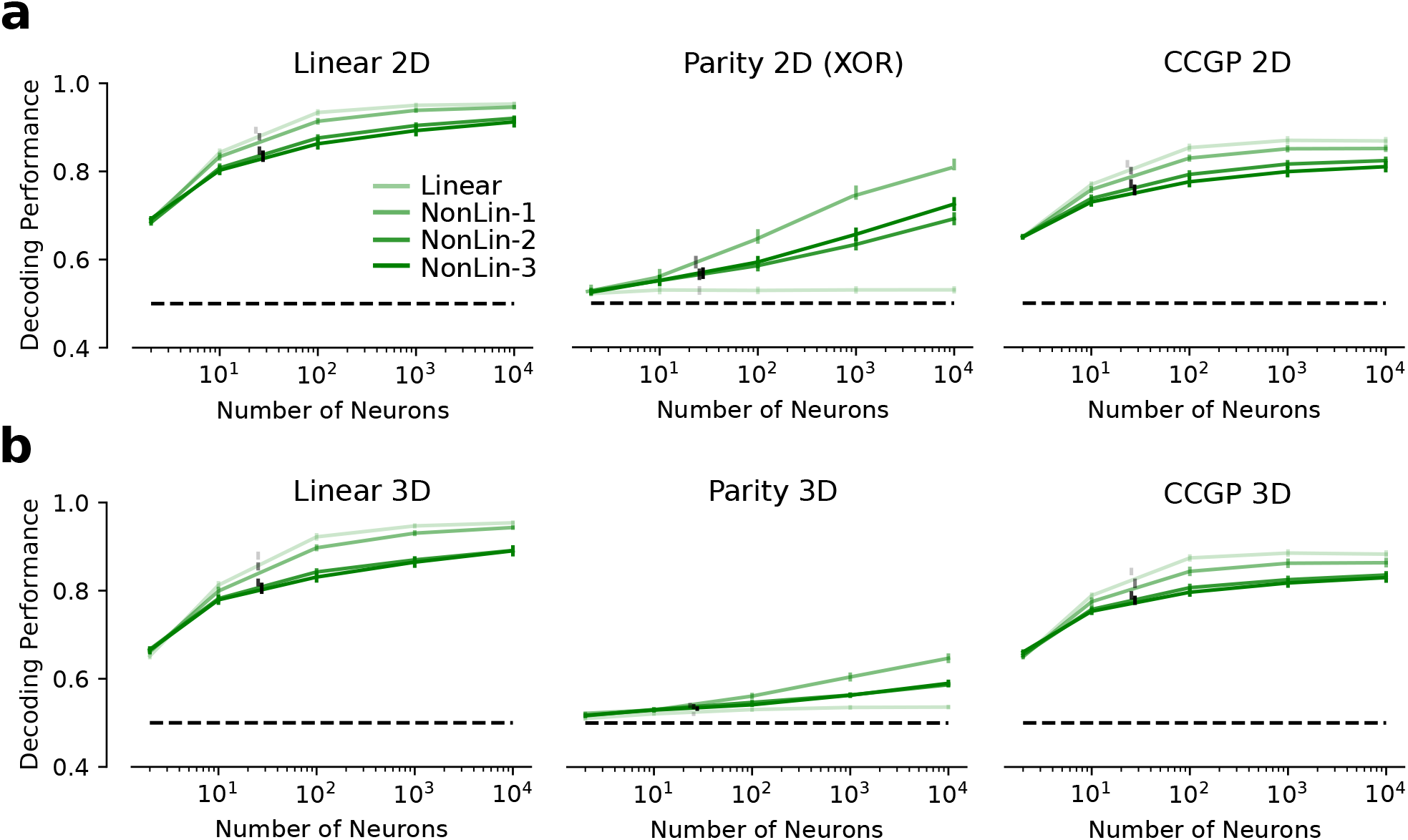
Performance on the easy (Linear) and complex (XOR) tasks as well as on the generalization benchmark (CCGP) grow as a function of the size of the surrogate population of neurons. (**a**) (left) Decoding Performance of a linear classifier trained to perform the easy task (Linear 2D) by reading out surrogate activity generated with the different encoding models (green). Surrogate activity generated with the linear encoding model produces the highest performance. Larger surrogate populations produce higher performances. Performance for the mean number of simultaneously recorded neurons across sessions (25.4 neurons) for the different encoding models is shown as black dots (horizontal jitter added on each value for visualization purposes). (middle) Same as (left) when the linear classifier was trained to perform the complex task (XOR). Surrogate activity generated with the linear encoding model was at chance level for all population sizes. Surrogate activity generated with an encoding model with only 1 hidden layer (NonLin-1), produced the highest decoding performance. (right) Same as (left, middle) for the generalization benchmark (CCGP, see Methods). Similar to the Linear 2D, surrogate activity generated with the linear encoding model produced the highest performance. In Fig. 7, the results for *N* = 100 are shown. For each session, task and number of neurons, the reported performance corresponds to the mean across the three possible pairs of whisker contacts (*C*_1_,*C*_2_), (*C*_1_,*C*_3_), and (*C*_2_,*C*_3_). (**b**) Same as (a) for the easy and complex tasks and generalization benchmark defined from contacts of the three whiskers simultaneously (*C*_1_,*C*_2_,*C*_3_) (see Methods). Results are qualitatively equivalent to (a). Errorbars in all panels correspond to s.e.m. across recording sessions.

**Figure S15:**
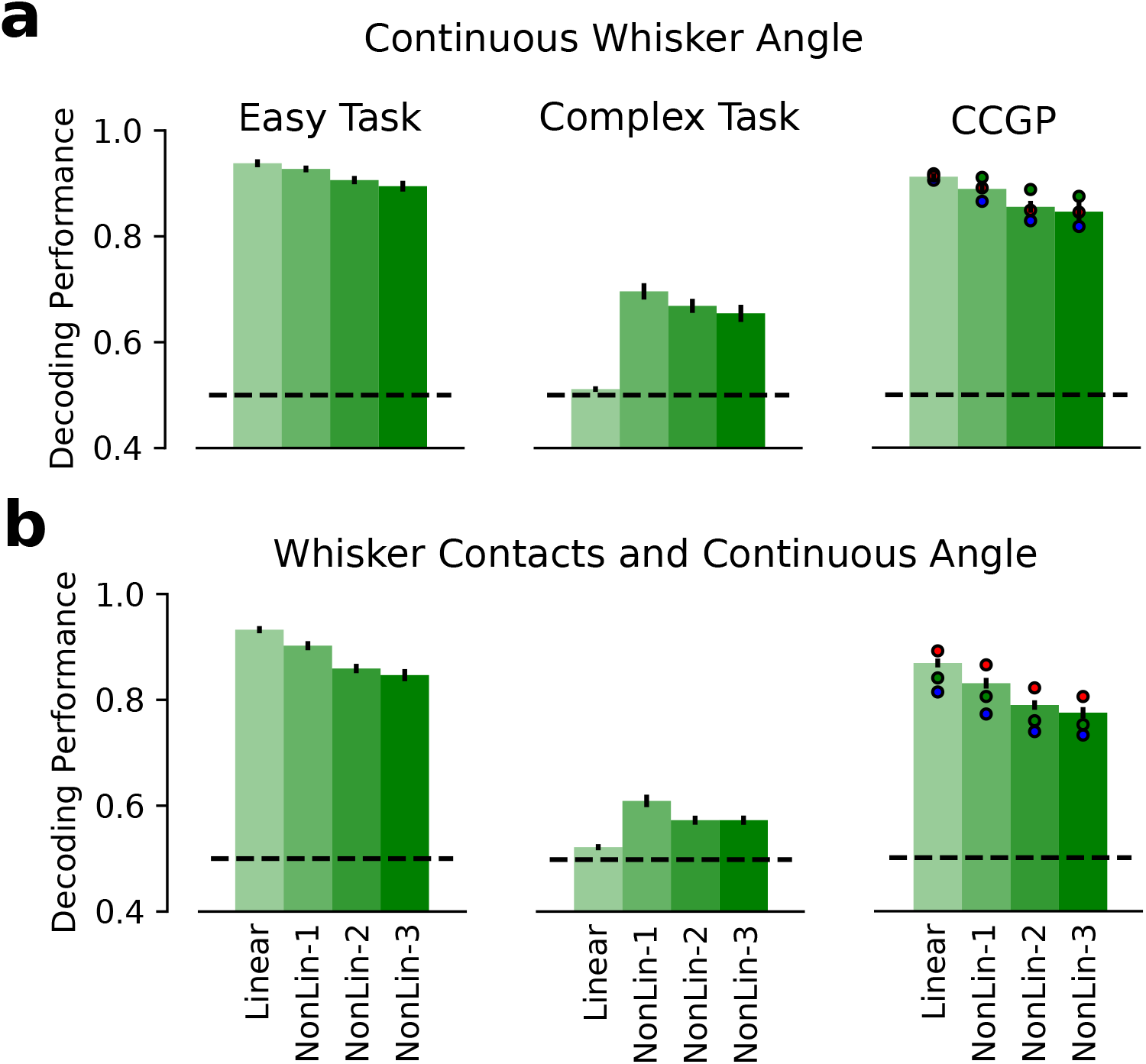
Qualitatively equivalent results to Fig. 7 were found when the easy and complex tasks and the generalization benchmark (CCGP) were defined with respect to the task variable continuous whisker angular position (a) and the interaction between the task variables whisker contacts and whisker continuous angular position (b) (see Methods). Results are shown for populations of 100 surrogate neurons. As in Fig. 7, errorbars in all panels correspond to s.e.m. across recording sessions.

**Figure S16:**
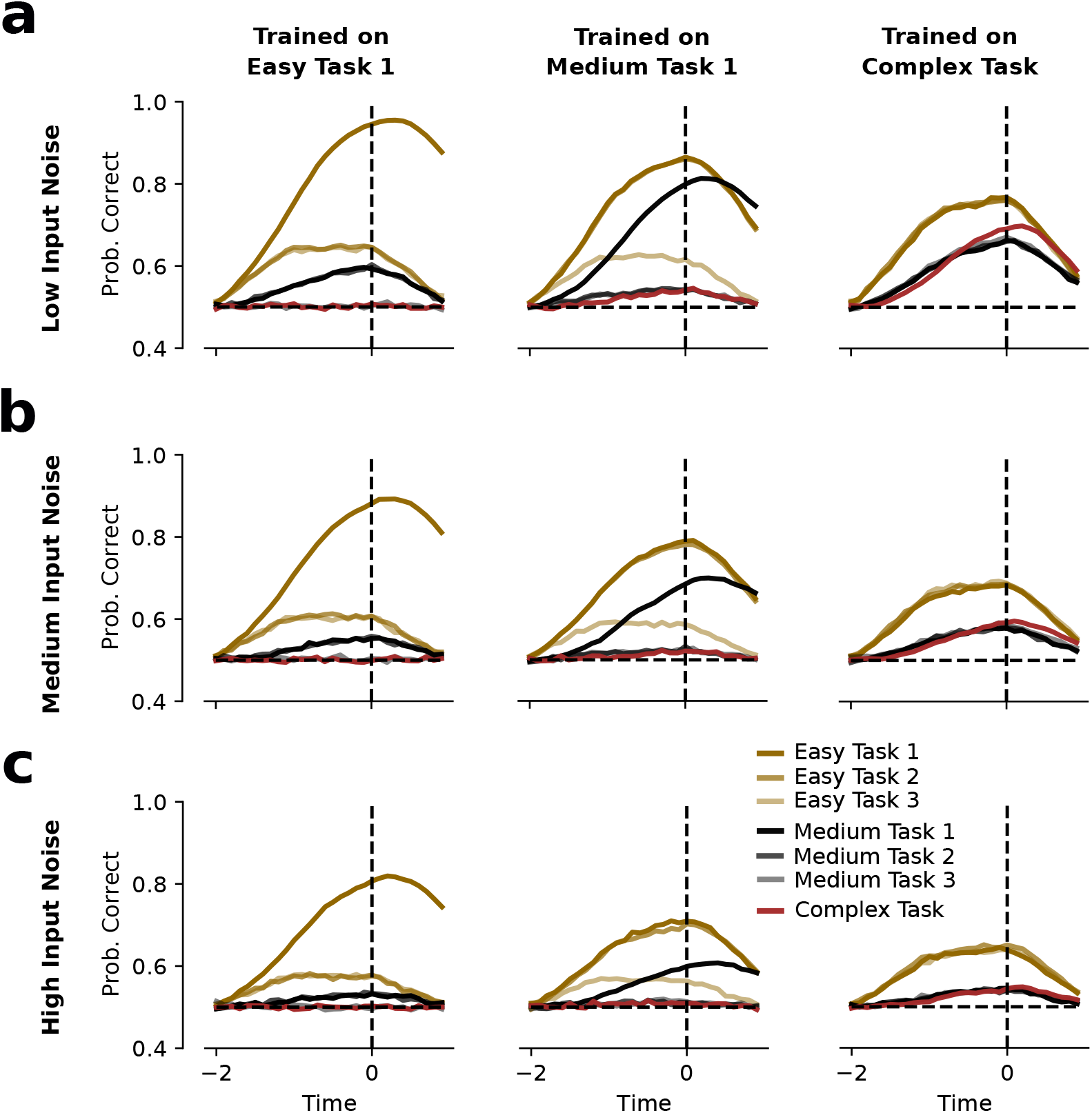
RNNs trained on a complex task produce the best trade-off between generalization and discrimination across all tasks and noise levels. (**a**) Probability of correct response (y-axis) as a function of time (x-axis) when the RNN was trained on the easy task 1 (left panel), the middle task 1 (central panel) and the complex task (right panel). RNNs were trained on low input noise (λ_*low*_ = 0.23 and λ_*high*_ = 0.77). For each network, additional readout weights on the activity of the artificial units were trained to perform the rest of the tasks (see Methods). While an RNN trained on easy task 1 produced the best performance for easy task 1, the neuronal representations were not well suited for the rest of tasks (left). When an RNN was trained on the middle 1 (central) and complex (right) tasks, it produced representations that allowed the performance of many different tasks. This came at the expense of losing performance for the easy task 1. (**b-c**) The same qualitative results were obtained when medium (b; λ_*low*_ = 0.3 and λ_*high*_ = 0.7) and high (c; λ_*low*_ = 0.35 and λ_*high*_ = 0.65) noise levels were used instead. For each panel the performance curves correspond to the mean across 50 random realizations of input patterns and tasks (see Methods).

**Figure S17:**
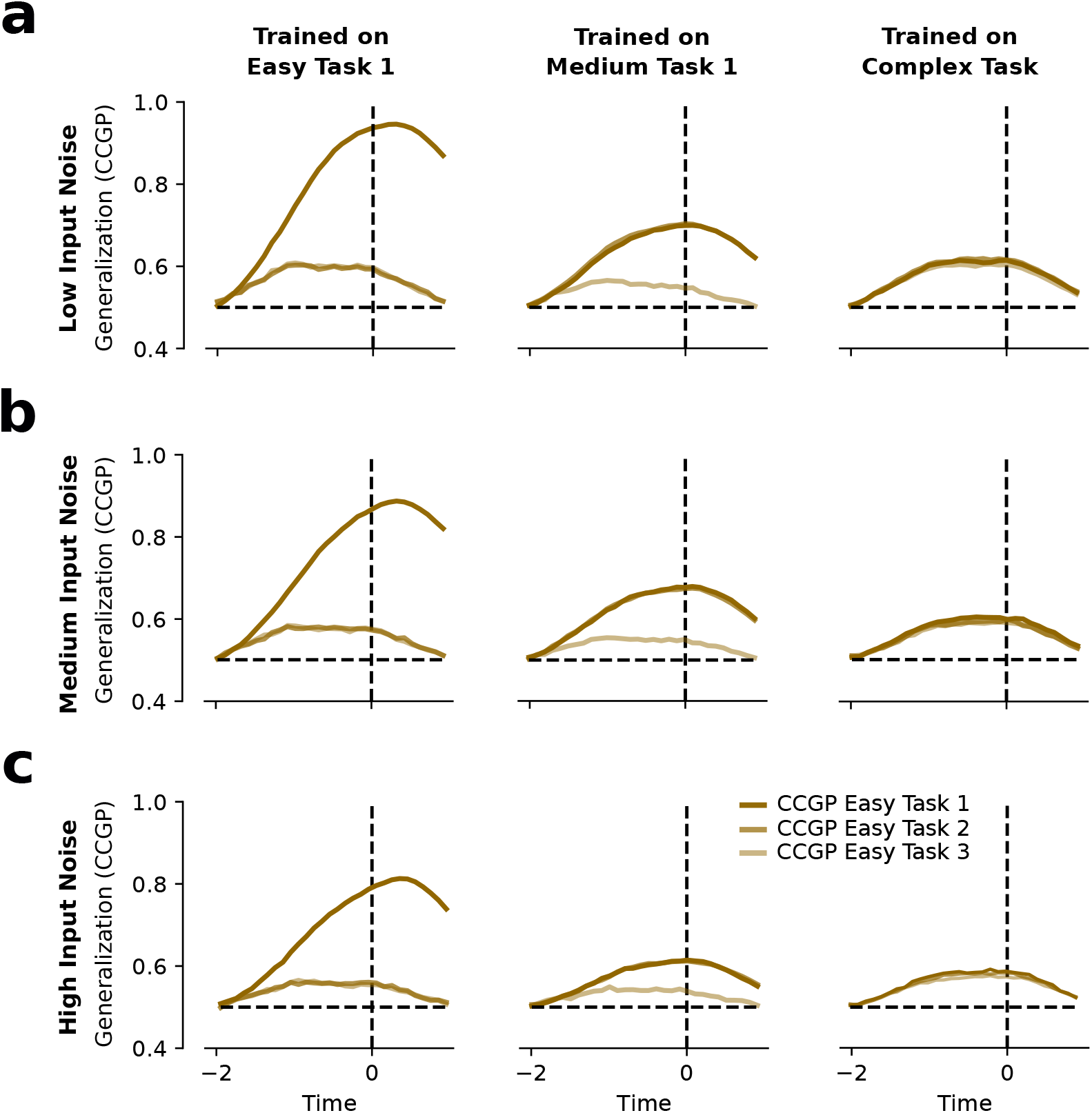
Generalization performances (CCGP) are above chance for even the RNNs trained on the complex task. (**a**) Generalization performance defined as cross-condition generalization performance (CCGP; y-axis) as a function of time (x-axis) when the RNN was trained on the easy task 1 (left panel), the middle task 1 (central panel) and the complex task (right panel). RNNs trained on low input noise (λ_*low*_ = 0.23 and λ_*high*_ = 0.77). While abstraction (CCGP) is high for easy task 1 when the RNN was trained on the easy task 1, it is low for easy tasks 2 and 3. Since medium task 1 is defined as the 2D-XOR between C1 and C2, CCGP was higher for easy task 1 and 2 (C1 and C2) than for easy task 3 (C3). For the RNN trained on the complex task, CCGP was significantly above chance for all easy-task variables. (**b-c**) The same qualitative results were obtained when medium (b; λ_*low*_ = 0.3 and λ_*high*_ = 0.7) and high (c; λ_*low*_ = 0.35 and λ_*high*_ = 0.65) noise levels were used instead. Results in all panels are qualitatively similar to Supplementary Fig. S16. For each panel the performance curves correspond to the mean across 50 random realizations of input patterns and tasks (see Methods).

**Figure S18:**
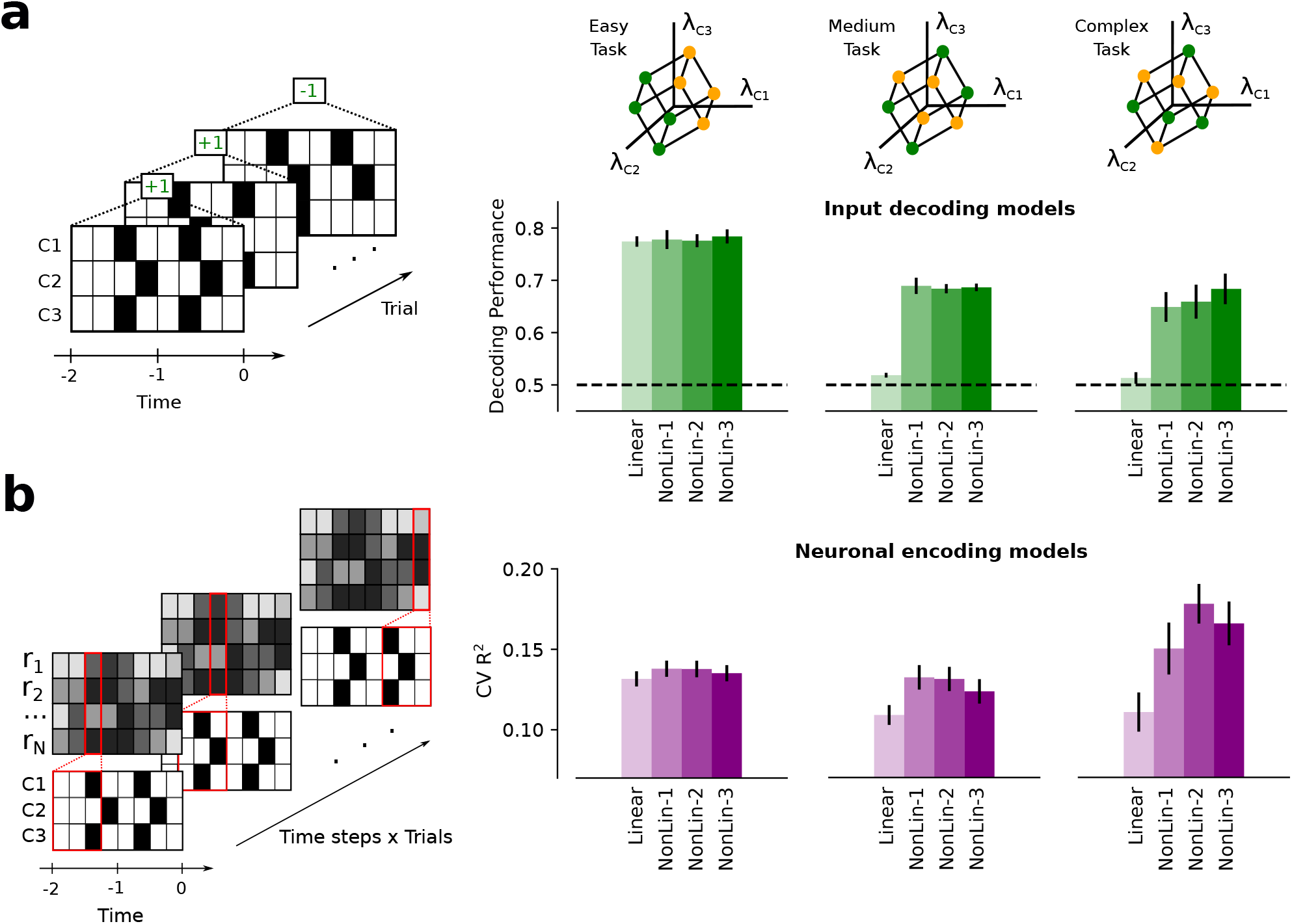
The advantage of the non-linear encoding models grows with the difficulty of the integration task. (**a**) Similar to Fig. 3d, linear and non-linear classification models that read out from the input space (x-axis), were trained to perform the easy (left panel), medium (central panel) and complex tasks (right panel). On the easy task, both linear and non-linear models performed equally well, as shown by decoding performances (y-axis) of the different models. On the contrary, only non-linear classifiers that allow for complex cue combination, performed above chance on the medium (central) and complex (right) tasks. The behavioral results obtained on the whisker-based discrimination task (see Fig. 3) are aligned with the easy task (left panel). In all panels errorbars correspond to the s.e.m. across four different network realizations. (**b**) Similar to Fig. 4d, cross-validated (CV) *R*^2^ on explaining artificial units activity (y-axis), is plotted against the different encoding models (x-axis). RNNs trained to perform tasks that require non-linear integration of sensory cues are better explained by an encoding model with non-linear mixed selectivity (central and right panels), while a non-linear encoding scheme provide marginal additional explanatory power for the easy task RNN (left panel). The higher the complexity of the trained task, the higher the advantage of non-linear encoding models on explaining the activity of the artificial units. In contrast to (a), the encoding properties of S1 are aligned with those of RNNs trained to perform tasks that require non-linear combination of sensory evidence (see Fig. 4). In all panels errorbars correspond to the s.e.m. across 240 artificial units (4 networks × 60 units).

